# Expanding the diversity of nitroxide-based paramagnetic probes conjugated to non-canonical amino acids for SDSL-EPR applications

**DOI:** 10.1101/2024.12.18.629154

**Authors:** Maxime Bizet, Áron Balázsi, Frédéric Biaso, Deborah Byrne, Emilien Etienne, Bruno Guigliarelli, Philippe Urban, Pierre Dorlet, Gilles Truan, Guillaume Gerbaud, Tamás Kálai, Marlène Martinho

## Abstract

Understanding protein structure requires studying its dynamics, which is critical to elucidating its functional role. Over time, biophysical techniques have revolutionized this field, offering remarkable insights into the structure-function relationship. Among these, Site-Directed Spin Labelling (SDSL) combined with Electron Paramagnetic Resonance (EPR) is a powerful method delivering structural data at the residue level, irrespective of protein size or environment. Traditional nitroxide labels, which target cysteine residues, often face limitations when these residues are essential for protein structure or function. To address this, alternatives have been proposed as the use of non-canonical amino acids (ncaa) coupled with specific nitroxide labels. This study introduces ^14^N-HO-5223, a novel nitroxide label specific to the *p*AzPhe ncaa, alongside its ^15^N-derivative. These labels were grafted at two sites of the model protein, the diflavin Cytochrome P450 reductase. For comparative purpose, two already reported labels were also used. Continuous wave (cw) EPR spectroscopy validated the HO-5223 label as an effective reporter of protein dynamics. Additionally, Double Electron-Electron Resonance (DEER) measurements provided distance distributions between the semi-quinone FMNH^•^ state of the CPR and all nitroxide labels. These results expand the toolkit of the ncaa-nitroxide pairs, enabling EPR-based structural studies of proteins where cysteine modification is impractical, further advancing our ability to decode protein dynamics and function.

## Introduction

Site-Directed Spin Labelling combined with Electron Paramagnetic Resonance (SDSL-EPR) and Double Electron-Electron Resonance (DEER) is now established as a robust method to gain dynamics and structural insights respectively, into nucleic acids, soluble and membrane proteins, *in vitro* and *in cell*.^[1–5]^ This approach is based on the grafting of a spin probe, usually a nitroxide radical, typically on native or engineered cysteine residues.^[6–8]^ The power of the approach resides in the fact that the continuous wave (cw) EPR spectra of nitroxide labelled proteins are very sensitive to the local structural environment of the label and are modified accordingly.^[6,9,10]^ Protein-protein interactions, as well as conformational changes or induced folding can then be explored.^[11,12]^ Using DEER, distance distributions can be accessed as well, in a 1/r^3^ manner, when two spin probes are present within a single protein, or on two partner proteins, through spin-spin dipolar interaction measurements.^[13,14]^ However, in some cases, one or too many cysteines are present in the primary sequence, and are not ideally situated or too numerous for labelling, which makes their mutation a necessary step before labelling. Moreover, cysteine residues may play critical roles in binding sites, catalytic activity, or structural stabilization of the protein.^[15]^ Nitroxide labels reactive to tyrosine residues have been proposed as a promising alternative.^[15,16]^ However, they face similar limitations as those for cysteine residues, and the labelling conditions were found to be harsh for certain proteins.

In recent years, several approaches have been proposed offering alternatives to cysteine labelling such as *i)* nitroxides^[17,18]^ or *ii)* triarylmethyl radicals^[19]^ both reactive to non-canonical amino acids (ncaa), *iii)* metal centers.^[20–24]^ The two latter types of label were specifically designed for distance measurements, whereas nitroxide labels offer the added capability to investigate protein dynamics as well. Endogenous radicals, such as flavin cofactors, can be added to this panoply of paramagnetic centers, allowing distances measurements as well, although not many examples have been reported.^[17,25,26]^

Labelling reactions involving non-canonical amino acids are determined by the chemical nature of each ncaa.^[27]^ For spin labelling, one can essentially distinguish Cu(I)-assisted reactions (Cu(I)-catalyzed Azide-Alkyne Cycloaddition or CuAAC)^[28,29]^ *vs*. Cu-free click reactions (Strain-Promoted Azide-Alkyne Click reaction or SPAAC^[30,31]^; Strain-Promoted Inverse-Electron-Demand Diels–Alder Cycloaddition or SPIEDDAC^[32]^; Strain-Promoted Alkyne-Nitrone Cycloaddition or SPANC^[33]^ and Staudinger ligation^[34,35]^; oxime/hydrazone ligation^[36–40]^). In the case of CuAAC reaction, the presence of Cu(I) can be harmful to the biological system, even with a chelator present. More confidentially, Pd-catalyzed reaction has also been reported.^[41]^ Among all possible ncaa,^[42]^ SCO-(carbamate-coupled Strained Cyclic Octyne) or BCN (Bi-Cyclic Nonyne)-based lysine,^[43,44]^ PrK (propargyl lysine),^[43]^ *p*AcPhe (*p*-acetyl-L-phenylalanine),^[45,46]^ *p*AzPhe (*p*-azido-L-phenylalanine),^[47]^ *p*IPhe (*p*-iodo-L-phenylalanine),^[41]^ *p*ENPhe (*p*-ethynyl-L-phenylalanine),^[48]^ *p*PrPhe (*p*-propargyloxy-L-phenylalanine),^[48]^ *p*2yneY (*p*-O-pentynyl-L-tyrosine)^[48]^ have been reported as targets for spin labelling (noted hereafter as ncaa*, star standing for the grafted label).

Recently, we reported cw EPR experiments as well as DEER experiments on semi-quinone flavine/nitroxide pairs in the Cytochrome P450 Reductase (CPR) protein (soluble His-Tag-Δ61-CPR).^[17]^ CPR is a flavoprotein located in the cytoplasmic side of the endoplasmic reticulum. CPR accepts electrons from the nicotinamide adenine dinucleotide phosphate (NADPH), which then transit via CPR’s flavin cofactors (from flavin adenine dinucleotide (FAD) to flavin mononucleotide (FMN)), reaching one of its physiological final electron acceptors, the heme iron of Cytochrome P450 (CYP). This electron transfer (ET) requires domain movements between the so-called “locked/closed” (for intra-ET “FAD to FMN”) and “unlocked/open” (for inter-ET “FMN to the heme iron of CYPs”) states. We demonstrated that these two states exist in solution when CPR is in its specific redox resting, air stabilized state, known as the FAD/FMNH^•^ state. Additionally, we identified a more widely open electrostatic driven state, compared to the “unlocked/open” reported one, which we interpreted as an efficiency conformation for interactions with external partners, such as Cytochrome c or CYPs.

In this study, we used the previously mentioned CPR as a model system for further FMNH^•^-NO^•^ distance measurements. We compared an acid-catalyzed reaction involving the ncaa *p*AcPhe grafted with ^14^N-specific label HO-4120 (3-aminooxymethyl-2,2,5,5-tetramethyl-2,5-dihydro-1H-pyrrol-1-yl oxylradical),^[45]^ with a SPAAC-based reactions using *p*AzPhe, which was labelled with HO-4451 (1-Fluoro-N-[(1-oxyl-2,2,5,5-tetramethyl-2,5-dihydro-1H-pyrrol-3-yl)methyl]cyclooct-2-ynecarboxamide). ^[17,47]^ Additionally, we expanded labelling options by introducing a novel label, ^14^N/^15^N-HO-5223 ((((1R,8S,9s)-bicyclo[6.1.0]non-4-yn-9-ylmethyl 2,2,5,5-tetramethyl-2,5-dihydro-1H-pyrrole-3-carboxylate)-1-yl)oxydanyl), specifically designed for the ncaa *p*AzPhe (Figure 1). The ^15^N isotopic version of this label will enable advanced applications in studying protein-protein interactions and bio-orthogonal labelling. The synthesis of HO-5223 label proved to be simpler than that of the HO-4120 label. Additionally, the oxime/hydrazone ligation formed with the HO-4120 label required harsh conditions for biological systems, such as acidic pH. Although the use of a catalyst allows the reaction to proceed at a neutral pH, it still necessitates a reaction time of three days at room temperature.^[45]^ Our comparison will focus on several key parameters: i) labelling yields, ii) impact on protein activities, iii) geometry of the label including size, flexibility, and linkage to the ncaa, iv) capacity of the novel label to report on protein dynamics in cw EPR experiments and v) distance distributions observed in DEER experiments. We demonstrated that all the ncaa/nitroxide pairs presented in this study were effective reporters for investigating protein dynamics, regardless of their size or linkage. DEER experiments provided structural insights into CPR. As previously reported, these experiments confirmed the existence of two primary states of the one-electron-reduced CPR in solution. Finally, this work also opens up new opportunities for application in systems where cysteine cannot be modified, thereby expanding the toolkit available for studying protein dynamics.

**Figure 1.**
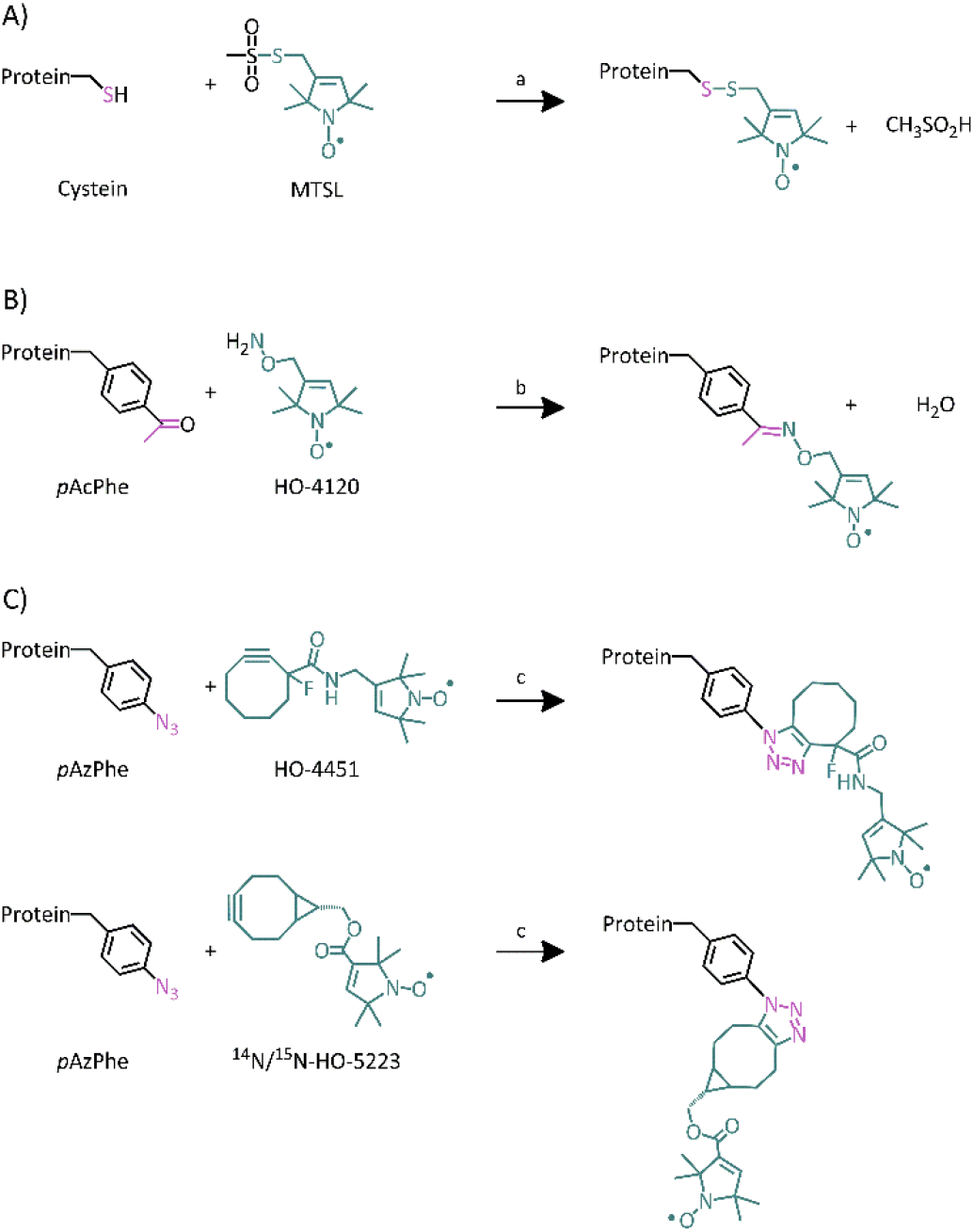
Ncaas, specific spin labels, and general labelling reaction schemes for A) cysteine with MTSL; B) aldehyde/alkoxyamine reaction for *p*AcPhe with HO-4120; C) SPAAC reaction for *p*AzPhe ncaa with HO-4451 and ^14^N/^15^N-HO-5223. Conditions: a) standard labelling conditions as previously reported^[6,14]^; b) rt, 72h, *p*-Methoxyaniline; c) rt, overnight, under dark conditions, Tris HCl 20 mM pH 7.4. See SI for details.

## Results and Discussion

### Synthesis and cw EPR characterization of spin label ^14^ N/^15^ N-HO-5223

The synthesis of the HO-4120 nitroxide label, designed to react specifically with the ncaa *p*AcPhe, has been reported previously.^[45]^ For labelling of the ncaa *p*AzPhe, the synthesis of HO-4451 spin label has also been reported.^[47]^ We have previously demonstrated its enhanced value in the study of CPR.^[17]^ Both ^14^N/^15^N-HO-5223 (Figure 1) have been synthesized through a two-step process, as depicted in scheme 1 (see SI for details). The ^15^N-HO-5223 label was specifically designed to provide a distinct EPR spectral signature, enabling further applications, particularly in studying protein-protein interactions.^[49]^ Although ^15^N-nitroxide labels have already been reported since the 1980s,^[50–53]^ none have been applied to the study of biological systems in presence of ncaas. The limited diversity of EPR signatures (three-line spectra) had historically hindered the simultaneous analysis of two sites within a single protein or between two interacting partner proteins. Previously, we reported a cysteine-specific β-phosphorylated nitroxide (referred to as PP), with a phosphorus atom near the NO group.^[49]^ This label exhibited a six-line spectrum EPR at room temperature and was successfully combined with a conventional nitroxide label (maleimido proxyl, displaying a 3-line EPR spectrum) to monitor induced folding in a model Intrinsically Disorded Protein (IDP) in the presence of its partner.^[54]^ However, the superposition of these signatures resulted in a complex spectrum, which could be simplified by employing a ^15^N isotope.

Reaction scheme for the synthesis of ^14^N/^15^N-HO-5223 label. Conditions: a) SOCl_2_, Pyr, toluene, 0°C to rt, 1h; b) Et_3_ N, DMAP, DCM, rt, 12h.^[55]^ See SI for details.

**Scheme 1.**
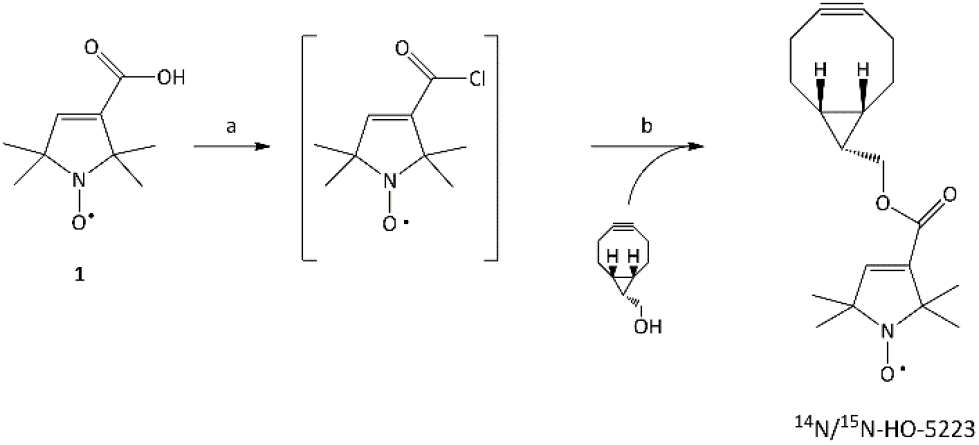
Reaction scheme for the synthesis of ^14^N/^15^N-HO-5223 label. Conditions: a) SOCl_2_, Pyr, toluene, 0°C to rt, 1h; b) Et_3_ N, DMAP, DCM, rt, 12h.^[55]^ See SI for details.

Unbound in solution, all ^14^N labels (HO-4120,^[46]^ HO-4451 and HO-5223) displayed typical, highly mobile three-line cw EPR spectra (Figures 2 and S1). These spectra arose from the hyperfine interaction (A_iso_(^14^N) ≈ 1.5 mT) between the electronic magnetic moment of the unpaired electron and the nuclear magnetic moment of the ^14^N atom (I(^14^N) = 1).^[6,56]^ For ^15^N-HO-5223 nitroxide label, the cw EPR spectrum showed a 2-line pattern (Figures 2 and S2) centered at g_iso_ = 2.008 with a A_iso_(^15^N) ≈ 2.2 mT, which is 1.4 times the A_iso_(^14^N) value. This was consistent with expectations and aligned with previously reported results for other ^15^N-nitroxides (I(^15^N) = 1/2).^[51,52,57,58]^ As for the case of ^14^N-nitroxide labels, changes in the cw EPR spectra of the ^15^N-HO-5223 label, depending on the rotational correlation time, provided insights into its mobility. This, in turn, offers information about its local structural environment (Figures S1 and S2). The ^15^N-HO-5223 label could then be used, alone or in combination with a ^14^N-nitroxide, to report on protein dynamics within a single protein or investigate protein-protein interaction.

**Figure 2.**
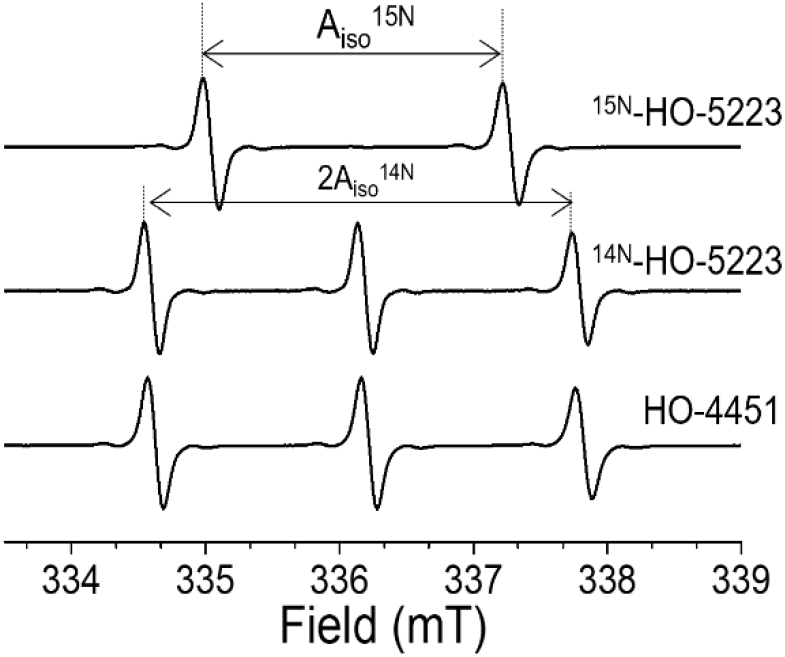
Cw EPR spectra of nitroxide labels recorded at room temperature. Conditions: [label] = 100 µM in Tris HCl 20 mM pH 7.4, P = 10 mW, MA = 0.1 G, *v* = 9.45 GHz.

### Incorporating ncAAs in soluble CPR

In order to graft nitroxide labels at selected positions, two non-canonical amino acids, *p*AcPhe or *p*AzPhe, were incorporated into *Homo sapiens* CPR (His-Tag-Δ61 wt, separately numbering based on the NCBI Reference Sequence NP_001382342, see SI) at positions 157 (FMN domain) and 668 (FAD domain), noted hereafter as Q157^*p*AzPhe^, Q157^*p*AcPhe^, K668^*p*AzPhe^ or K668^*p*AcPhe^ (Figures 3, S3 to S6 and details in SI). Overall, incorporation of ncaa resulted in good yields of protein production, with the exception of Q157^*p*AzPhe^, when, compared to other samples, only about one-third of the protein was obtained (Table S1). The optimization of the ncaa position and choice is often not straightforward and typically relies on a trial-and-error approach, but optimal results were obtained when the position is on the solvent exposed surface. This highlights the importance of expanding the variety of ncaas together with specific labels used, as such diversity is crucial for improving labelling efficiency and enhancing the success of experiments.

**Figure 3.**
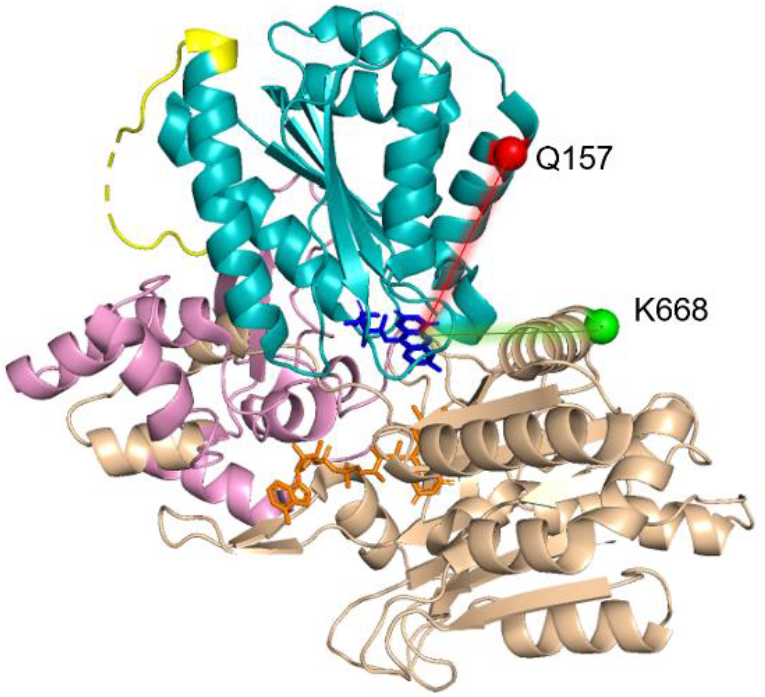
Q157 and K668 residues highlighted as red and green spheres respectively in the “Locked/closed” state crystal structure of soluble *Homo sapiens* CPR (pdb. 5fa6). The FAD and FMN domains are represented in brown and cyan ribbons respectively, FAD and FMN co-factors are shown as sticks in orange and dark blue, linker is shown as light pink ribbon, the flexible loop is shown in yellow. Red and green lines represent distances between FMNH^•^ and the label at positions Q157 and K668, respectively.

### Conformational study of nitroxide labels

Labels HO-4120, HO-4451, ^14^N- or ^15^N-HO-5223 had each a distinct structure and could be grafted to either *p*AcPhe (for HO-4120) or *p*AzPhe (Figure 1). Notably, the linkers connecting the ncaa to the nitroxide moiety vary in length and flexibility. A conformational analysis by quantum chemistry methods indicated that these linkers were all longer than the one in the cysteine-MTSL pair (Table 1 and figure 4). Yet, each label produced a similar number of rotamers to those generated by MTSL (Table 1). Additionally, the free rotation along the linker bonds (Figure 4) allowed the label to adopt conformations with either extended or more compact geometries (Figure S7). Note that a rigid cyclooctene structure was formed in conjunction with a triazole ring for the HO-4451 and HO-5223 labels when paired with *p*AzPhe. Consequently, the NO^•^-C_α_ distance varied for each ncaa/nitroxide pair (Figure 4 and table 1), with the larger distance variation for HO-4451 (6.00 Å) and the lowest one for HO-5223 (2.90 Å).

**Table 1.** Calculated^[a]^ N_NO_-C_α_^[b]^ distances (in Å) for grafted ncaa with nitroxide labels.^£^.

| Grafted aa | Cysteine | pAcPhe | pAzPhe |  |
| --- | --- | --- | --- | --- |
| Label | MTSL | HO-4120 | HO-4451 | HO-5223 |
| Number of rotamers | 23 | 23 | 18 | 17 |
| Minimum $N_{NO}-C_{\alpha}^{[b]}$ distances | 5.1 | 10.0 | 8.80 | 14.0 |
| Maximum $N_{NO}-C_{\alpha}^{[b]}$ distances | 8.3 | 13.3 | 15.8 | 16.9 |
| Average $N_{NO}-C_{\alpha}^{[b]}$ distance | 7.4 | 11.3 | 14.1 | 16.0 |
| Distance variation | 3.20 | 3.30 | 6.00 | 2.90 |

[a] see SI for computational details. [b] in Å. £ : MTSL linked to cysteine has been added for comparison.

**Figure 4.**
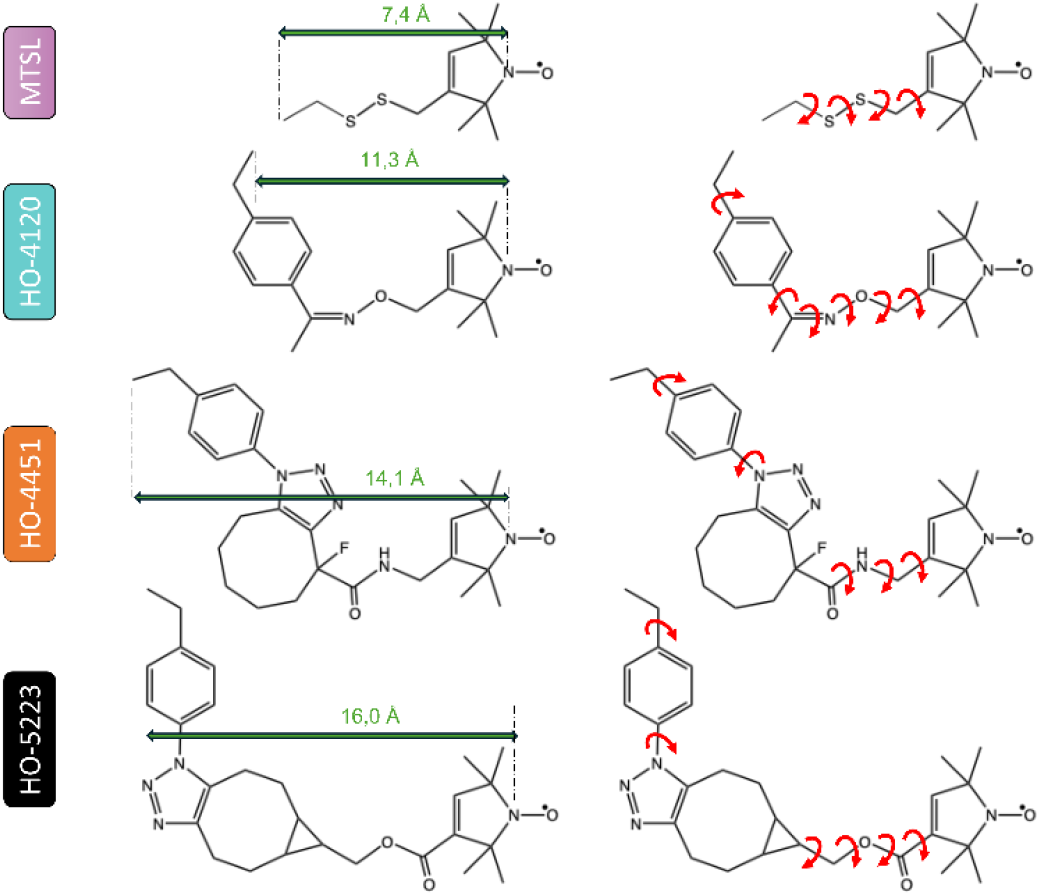
Comparative chemical structures of nitroxide spin labels grafted to ncaas used in this study. MTSL linked to cysteine has been added for comparison. Green straight arrows indicate calculated distances between the N atom of the NO^•^ group and the Cα of the terminal methyl group (see Table 2 and SI). Red arrows indicate calculated possible rotations.

### Cw EPR of labelled protein containing a ncaa

Each single CPR mutant was labelled with the aforementioned ^14^N- or ^15^N-labels, and will be referred to, using the following nomenclature: mutant^ncaa/label^ (Table 2). As previously reported, activity assays demonstrated that neither the incorporation of *p*AzPhe nor its spin labelling affected the ET capacity of CPR.^[17]^ However, in the specific case of Q157^*p*AcPhe/HO-4120^, a significant decrease in activity was observed (Table S2). To understand if this loss of activity was due to the labelling reaction conditions or to the grafting of the probe, a control experiment was conducted under the same reaction conditions but without the label. The observed loss of activity confirmed that the reduction in CPR activity was due to the labelling reaction conditions rather than to the presence of the label itself. Table 2 presents the labelling yields obtained for each labelled position, depending on the ncaa, nitroxide label, and labelling conditions (see SI for details). The yields of the *p*AzPhe ncaa labelling varied from very low to very high, depending on several critical points: *i)* position and label, *ii)* sensitivity of the *p*AzPhe ncaa to UV-light,^[59,60]^ *iii)* sensitivity of the *p*AzPhe ncaa to reductant present during protein expression. In the two last cases, the resulting quantity of the N_3_ chemical group could be lowered. Even after optimization of production, purification and labelling steps to reach high labelling yields, some variation could subsist and could not be standardized. Notably, the K668^*p*AzPhe^ mutant labelled with the ^15^N isotopically label, *i*.*e*. ^15^N-HO-5223, showed significant lower yields. This result was unexpected, as the presence of the isotope should not impact the reaction efficiency. Overall, the labelling reactions were efficient, most mutants could be labelled and used for further studies.

**Table 2.**
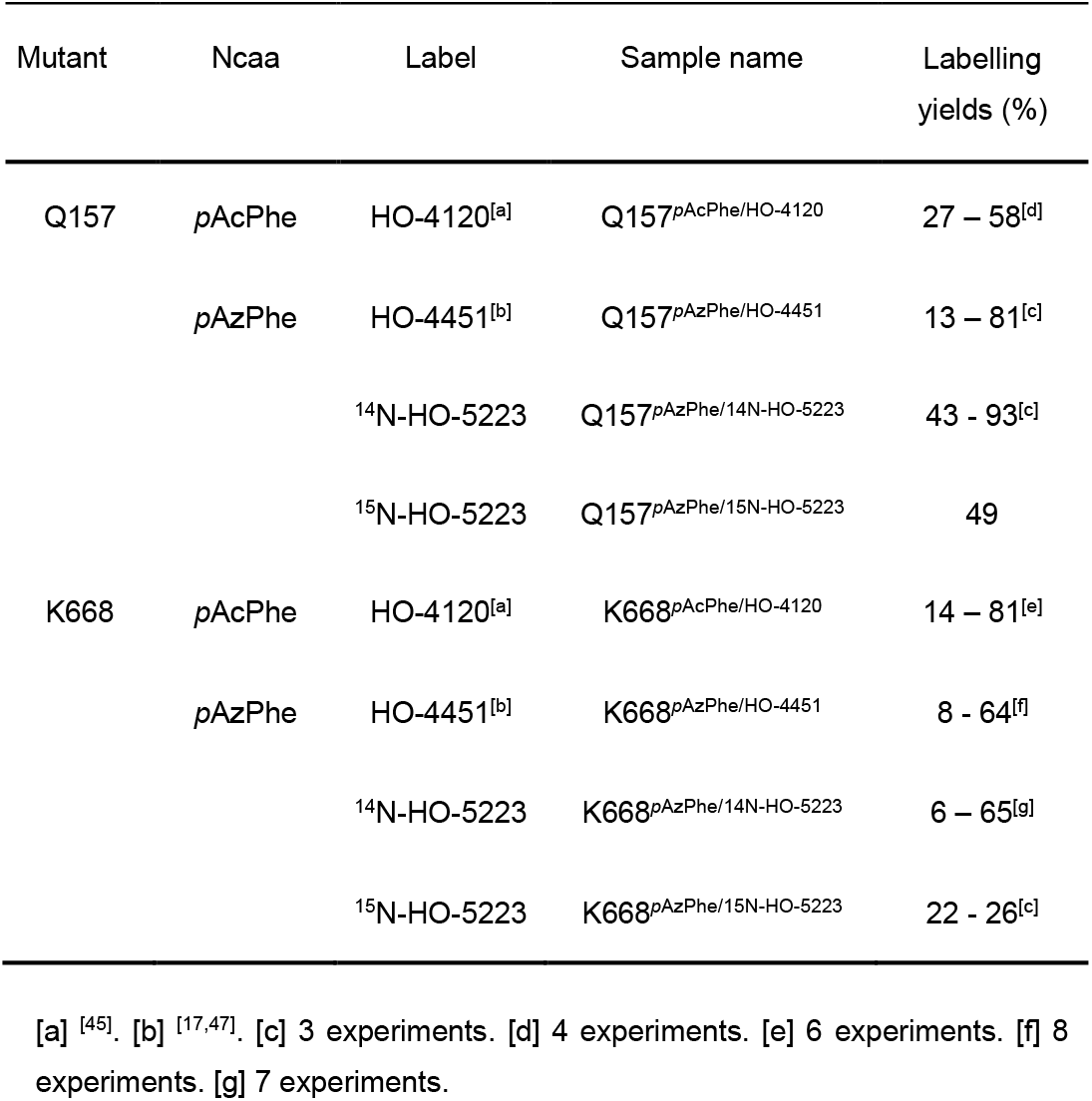
Nomenclature used for the different samples of labelled CPR mutants and labelling yields.

Cw EPR spectra of Q157_*p*AcPhe/HO-4120_ and K668_*p*AcPhe/HO-4120_ mutants are shown in figure 5. For both positions, the spectra were characteristic of a label in the so-called “intermediate regime of mobility”.^[11,61]^ Furthermore, the spectral shapes were similar, even if the K668^*p*AcPhe/HO-4120^ spectrum appeared slightly narrower, as evidenced by the more distinct hyperfine interaction with ^13^C atoms in the central line (marked with a blue star in figure 5). This similarity suggested that the label encountered a comparable local structural environment at both positions. HO-4120 has already been shown to be an effective reporter for local structure.^[45]^). As mentioned above, labelling of the *p*AcPhe ncaa required several hours reaction at pH 6 (See figure 1 and SI for details), conditions which were not optimal for the activity of the Q157^*p*AcPhe/HO-4120^ protein (Table S2). Consequently, we will focus on the results obtained using the *p*AzPhe ncaa.

**Figure 5.**
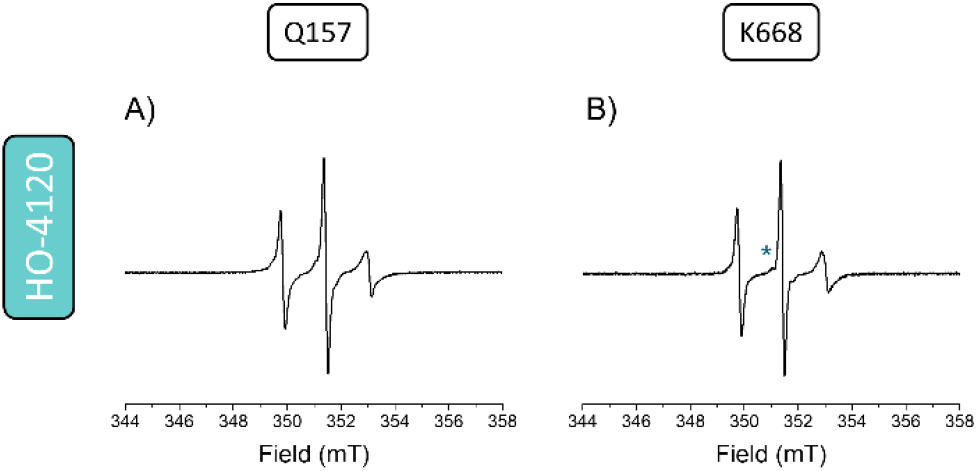
Room temperature cw X-band EPR spectra of labelled *Homo sapiens* CPR mutants. A) Q157_*p*AcPhe/HO-4120_ and B) K668_*p*AcPhe/HO-4120_. [Q157_*p*AcPhe/HO-4120_] = 55 μM, [K668^*p*AcPhe/HO-4120^] = 70 μM. Buffer: NaP 20mM pH 6 NaCl 500 mM. See SI for experimental conditions.

Figure 6 shows the cw EPR spectra for CPR Q157^*p*AzPhe^ and K668^*p*AzPhe^, labelled with the three labels: HO-4451, ^14^N- or ^15^N-HO-5223. When comparing the spectra for both positions labelled with HO-4451 (Figure 6, upper panel), the K668 position exhibited a slightly narrower spectrum, similar to the observation made with the HO-4120 nitroxide label on the *p*AcPhe ncaa (Figure 5). Additionally, a broader component was visible in the EPR spectrum of Q157^*p*AzPhe/HO-4451^ (indicated by black stars in the upper panel of figure 6). In the case of labelling with HO-5223, a supplemental and even more visible and broad signal was present in the Q157^*p*AzPhe/14N-HO-5223^ spectrum (blue arrows, middle panel in figure 6), with a split between the higher and lower field lines around 6.6 mT (indicated by the vertical blue dashed lines in figure 6). This broader line was not present on the K668^*p*AzPhe/14N-HO-5223^ spectrum. Since the same nitroxide label was grafted at both positions, the broad component observed in the Q157^*p*AzPhe/14N-HO-5223^ spectrum suggested the existence of an additional population where the label was in a more constrained local environment. This observation may indicate distinct protein conformations or label rotamers, similar to findings reported for the MTSL nitroxide.^[62]^ Another possibility was that the HO-5223 label, with its longer linker compared to HO-4120, could explore a broader and more distant environment, potentially accounting for the two distinct environments observed at positions Q157 and K668. Constantly, the spectrum of Q157^*p*AzPhe/15N-HO-5223^ displayed an overall lower flexibility compared to that of K668^*p*AzPhe/15N-HO-5223^, evidenced by a constrained spectral component indicated by blue arrows in the lower panel of figure 6. This component may arise from the same factors described above, with the distinctive spectral shape attributed to the ^15^N isotope (hyperfine interaction). These findings showed that despite using the same nitroxide label, the EPR spectra differed according to the targeted positions, highlighting the label sensitivity to local environments. Modifying the linker length of nitroxide labels could then serve as an additional tool for probing more distant environments. Overall, our results demonstrated that all three nitroxide labels used in this study effectively report on local environments, performing as robustly as conventional spin labels.

**Figure 6.**
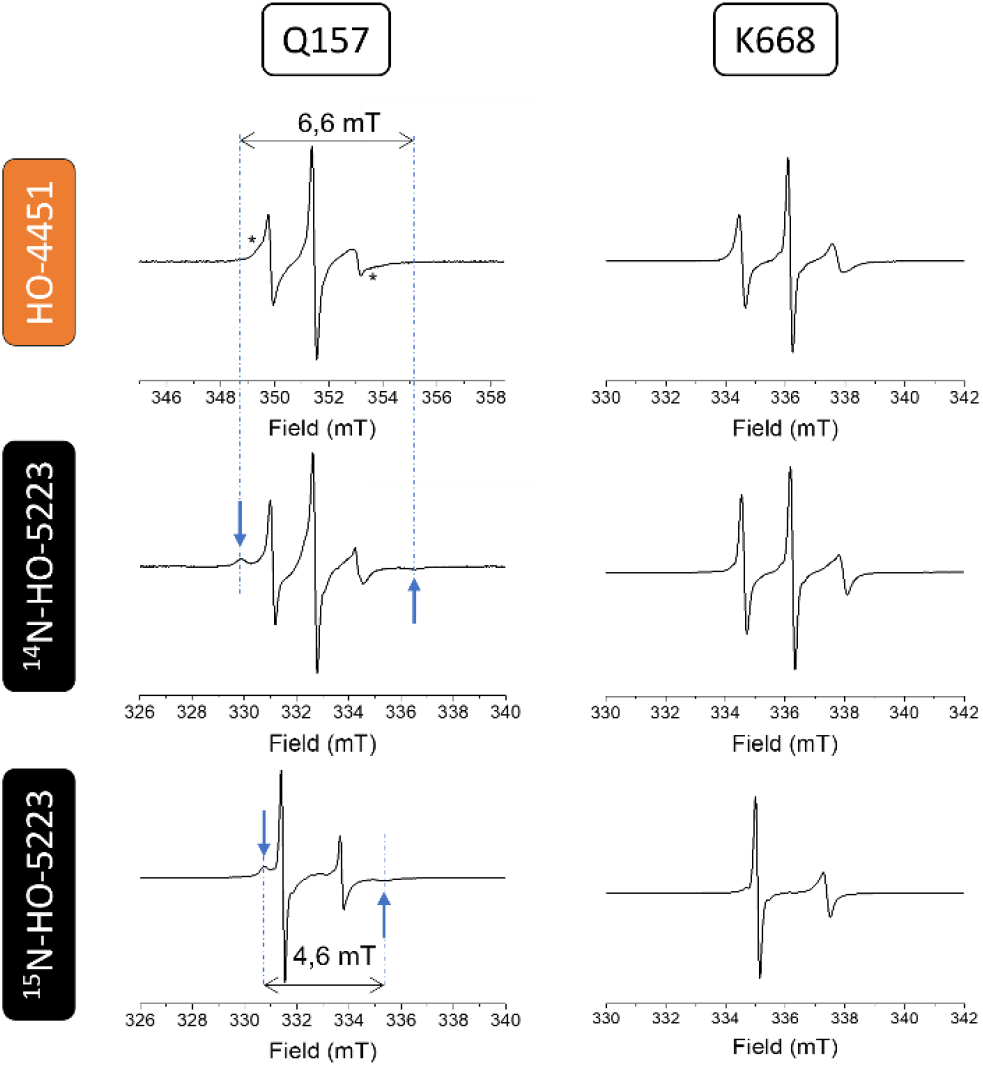
Room temperature cw X-band EPR spectra of labelled *Homo sapiens* CPR (left) Q157^*p*AzPhe^ and (right) K668^*p*AzPhe^ labelled with HO-4451(upper panel), ^14^N- (middle panel) and ^15^N-HO-5223 (lower panel). Arrows and stars indicate the broader component in the spectra. Vertical blue dashed lines are a guide for the eye. Protein concentration: 100 μM. Buffer: Tris HCl 20 mM pH 7.4. See SI for experimental conditions.

### Cw EPR characterization of FMNH* state in nitroxide labelled CPR

Adding 1 eq. of NADPH, the physiological electron provider for CPR, to the oxidized labelled CPR in air enabled us to prepare the FMNH^•^ semi-quinone state of the protein after a 10 minutes incubation (air stable species FAD/FMNH^•^, 1 electron reduced state)^[63–65]^ while simultaneously retaining the nitroxide label at positions Q157 or K668 (see SI).^[17]^ Cw EPR spectra were recorded in solution at room temperature for each sample (Figure 7).

**Figure 7.**
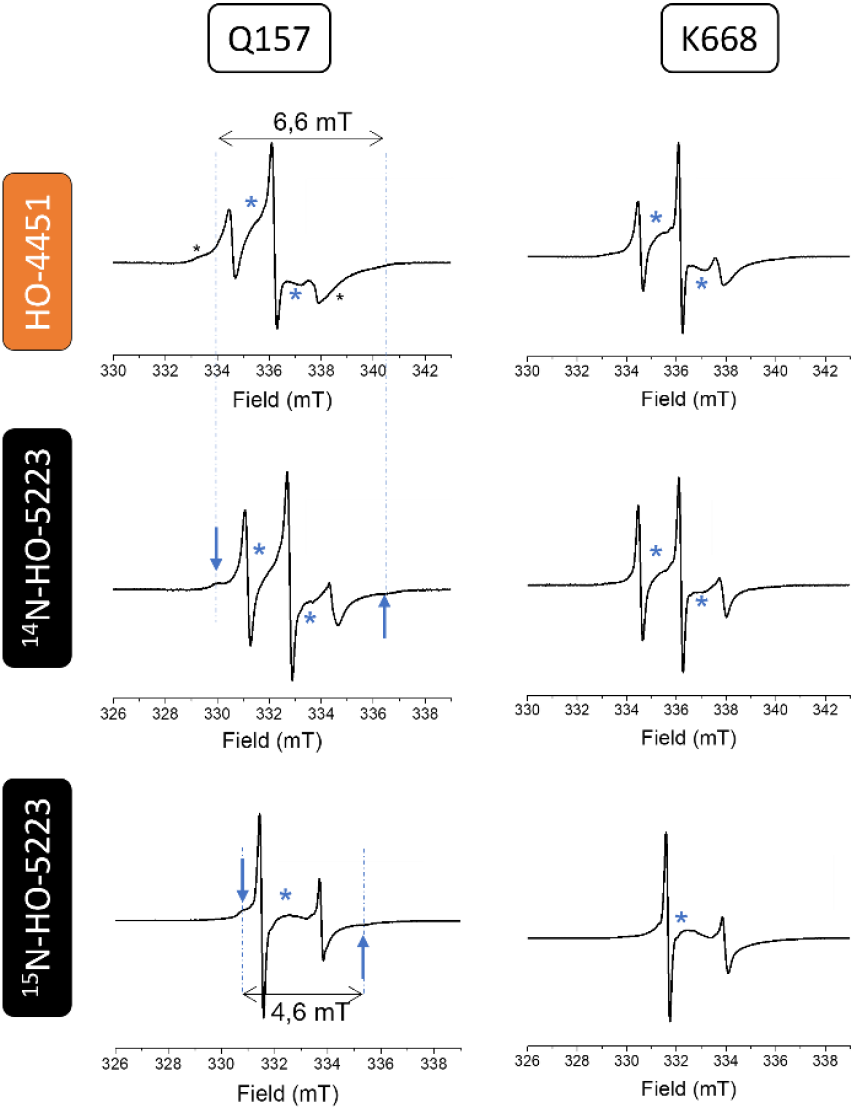
Cw X-band EPR spectra recorded at room temperature of labelled one-electron reduced *Homo sapiens* CPR mutants (left) Q157^*p*AzPhe^ and (right) K668^*p*AzPhe^ labelled with HO-4451(upper panel), ^14^N- (middle panel) and ^15^N-HO-5223 (lower panel). Blue stars indicate the FMNH^•^ signal. Black stars and blue arrows indicate the broader components in the spectra. Protein concentration: 100 μM. Buffer: Tris HCl 20 mM pH 7.4. See SI for experimental conditions.

As previously described, the FMNH^•^ EPR spectral shape was larger compared to the nitroxide one (∼3 mT). Both signals from the FMNH^•^ sq and the nitroxide label grafted to *p*AzPhe were present simultaneously for each position. Note that addition of NAPDH did not alter the spectral shape of nitroxide signal, indicating that the label dynamics remained unchanged in solution across both CPR redox states, *i*.*e*. oxidized and 1-electron reduced state, as previously described.^[17] 15^N-HO-5223 label displaying only a 2-line spectrum, the FMNH^•^ signal could be seen in the middle of the spectrum, as indicated by the blue star in figure 7. This label thus enabled the monitoring of both distinct, non-overlapping signals.

### Ncaa*-FMNH* distance measurements by DEER

Distance distributions between FMNH^•^ and nitroxide spin probes were investigated by DEER experiments, by varying the ionic strength of the solution using NaCl at different concentrations (0, 50 and 250 mM). As we previously reported, increasing ionic strength favored an extensively “unlocked/open” state in solution.^[17]^ The results obtained by DEER, using the three labels specific for *p*AzPhe ncaa, are shown in figures 8 and 9 for Q157 and K668 mutants respectively, together with MMM calculated FMNH^•^-NO^•^ distances (red curves). For all experiments, the pump (*v*_pump_) and observer (*v*_observe_) frequencies were chosen at the maximum of the FMNH^•^ and nitroxide signals on the echo field sweep (EFS), respectively (Figures S10 and S11). The observed modulation depth (λ) for DEER experiments on FMNH*/Q157_*p*AzPhe/HO-4451_ and FMNH*/K668_*p*AzPheHO-4451_ were consistent with our previous report, with λ values of 0.113 and 0.094 for position Q157 and K668 respectively. These values correspond to an approximative 50% spin labelling efficiency under our spectrometer configuration, as determined by the cw EPR spectra, regardless of the labelled position.^[17]^ For FMNH^•^/Q157^*p*AzPhe*^, a reduction in modulation depth was noted with ^14^N-HO-5223 (λ = 0.07) and was even more pronounced with _15_ N-HO-5223 (λ = 0.04), even with labelling yields of 50% as previously. Although resulting distance distributions were broader, consistent with the smoother oscillations, they remained centered at 28 Å in the absence of salt, as obtained using the HO-4451 label (Figure 10). This width of distance distribution may arise from the number of rotamers which the label can adopt, as predicted by the MMM^[66]^ software and by the good adequation with calculated distance distribution. We also examined the impact of ionic strength on modulation depth with HO-5223 labels. Upon addition of NaCl, no significant changes were observed in modulation depth (with λ values of 0.06 and 0.06 for FMNH^•^/Q157^*p*AzPhe/14N-HO-5223^ at 50 and 250 mM NaCl, respectively, and similarly, λ values of 0.03 and 0.03 for FMNH^•^/Q157^*p*AzPhe/15N-HO-5223^ at the same concentrations of protein). The time trace oscillations also remained unchanged, indicating stable FMNH^•^-NO^•^ distance distributions (Figure 8). This result was expected, as both probes are located within the same domain.

**Figure 8.**
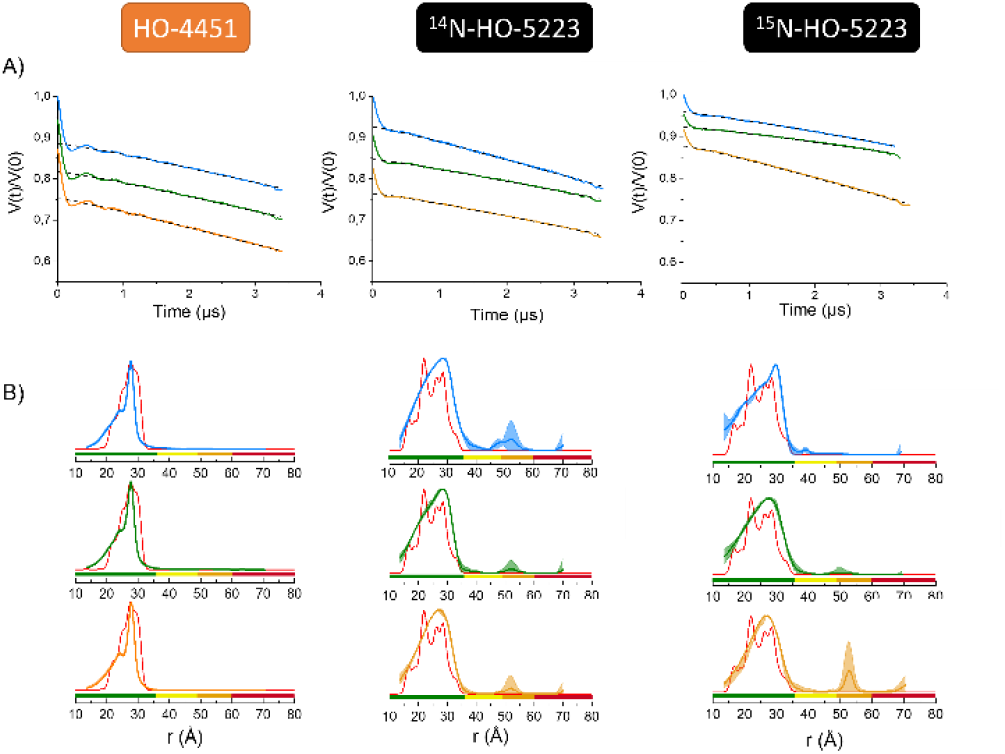
A) Experimental Q-band DEER traces recorded at 60 K for FMNH^•^/Q157^*p*AzPhe*^ in absence (blue line) or in presence of NaCl 50 mM (green line), 250 mM (orange line) using HO-4451 (left), ^14^N-HO-5223 (middle) and ^15^N-HO-5223 (right) labels. The black dashed lines indicate the baseline used for background correction. DEER traces were shifted vertically for the sake of clarity. B) Distance distributions obtained using the auto-computed mode in DeerAnalysis 2022^[67,68]^ (blue, green and orange lines) and calculated distance distributions for the *Homo sapiens* CPR “locked/closed” (red line) state (pdb. 5FA6 or 3QE2) obtained using the MMM software.^[66]^ For color code in the x axis, see SI. [Q157^*p*AzPhe*^] = 90 μM. Buffer: Tris HCl 20 mM pH 7.4, D_2_ O, 10% v/v *d*_*8*_-glycerol.

**Figure 9.**
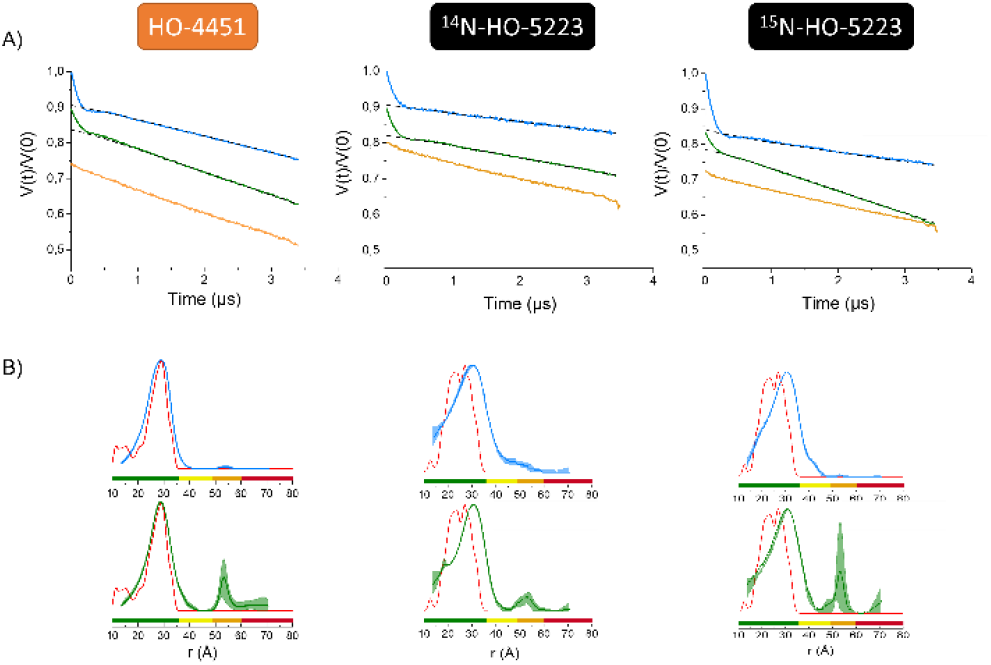
A) Experimental Q-band DEER traces recorded at 60 K for FMNH^•^/K668^*p*AzPhe*^ in the absence (blue line) or in the presence of NaCl 50 mM (green line), 250 mM (orange line) using HO-4451 (left), ^14^N-HO-5223 (middle) and ^15^N-HO-5223 (right) labels. The black dashed lines indicate the baseline used for background correction. DEER traces were shifted vertically for the sake of clarity. B) Distance distributions obtained using the auto-computed mode in DeerAnalysis 2022^[67,68]^ (blue, green and orange lines) and calculated distance distributions for the *Homo sapiens* CPR “locked/closed” (red line) state (pdb. 5FA6 or 3QE2) obtained using the MMM software.^[66]^ For color code in the x axis, see SI. [K668^*p*AzPhe*^] = 90 μM. Buffer: Tris HCl 20 mM pH 7.4, D_2_ O, 10% v/v *d*_*8*_-glycerol.

**Figure 10.**
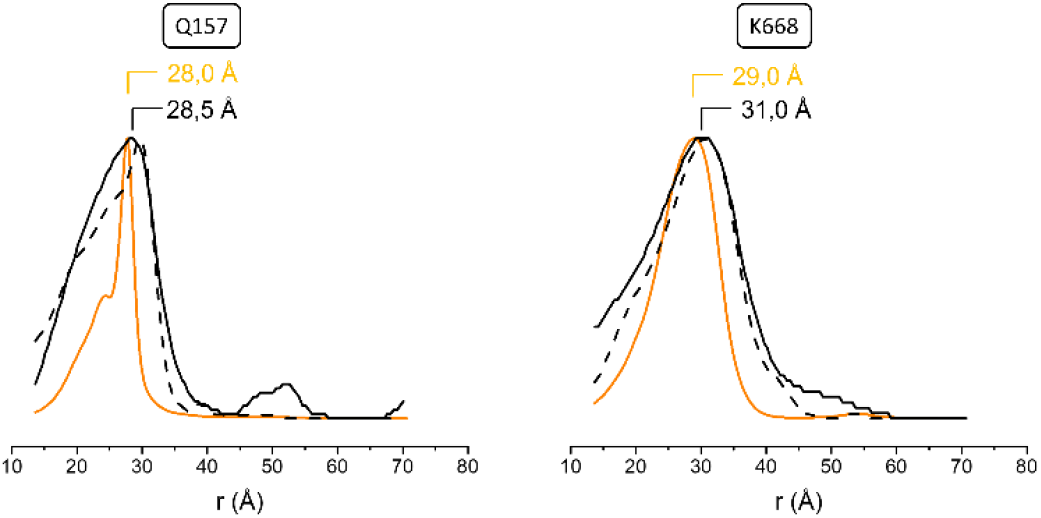
Comparison of distance distributions obtained using the auto-computed mode in DeerAnalysis 2022^[67,68]^ for the *Homo sapiens* FMNH^•^/Q157^pAzPhe*^ (left) and FMNH^•^/K668^*p*AzPhe*^ using HO-4451 (orange line), 14N-HO-5223 (black line) and 15N-HO-5223 (dashed black line) labels. [K668^*p*AzPhe*^] = [Q157^*p*AzPhe*^] = 90 μM. Buffer: Tris HCl 20 mM pH 7.4, D_2_ O, 10% v/v *d*_*8*_-glycerol.

For FMNH^•^/K668^*p*AzPhe/14N-HO-5223^, the modulation depth decreased less upon the addition of 50 mM of NaCl (λ reduced from 0.09 to 0.08) compared to FMNH^•^/K668^*p*AzPhe/15N-HO-5223^ (λ reduced from 0.15 to 0.05). However, at 250 mM of NaCl, the modulation depth was completely suppressed, suggesting a loss of measurable spin-spin dipolar interaction (Figure 9). As previously observed, the absence of modulation depth and oscillations indicated that the FMNH^•^-NO^•^ distance extended beyond the detectable range of pulsed EPR spectroscopy under our experimental conditions (> 8 nm was not detected in those conditions).^[17]^ Consequently, no distance distribution could be extracted, reinforcing the fact that the FMNH^•^-NO^•^ separation increased with high salt concentrations, favoring a more “unlocked/open” state of the one-electron reduced CPR form (FAD/FMNH^•^ state) in solution. As for the Q157 position, distance distributions measured with the HO-5223 labels were broader and centered at 31 Å, compared to 29 Å measured with the HO-4451 label (Figure 10).

Overall, the modulation depth in presence of increasing quantities of salt was consistent with the decrease of the dipolar interaction due to a longer distance between the flavin and the nitroxide centers. Nevertheless, although the labelling yields were similar for each position, the modulation depth decreased when using the HO-5223 label, and even more when using the 15N-HO-5223 label. This observation was unexpected and would need further experiments for a better understanding.

MMM^[66]^ (Multiscale Modelling of Macromolecules) software, an open-source modelling toolbox, was used on the crystal structure of soluble *Homo sapiens* CPR for the “locked/closed” (pdb. 5FA6) conformation^[69]^ and on the *Saccharomyces cerevisiae/Homo sapiens* CPR for the “unlocked/open” chimeric conformation (pdb. 3FJO).^[70]^ Spin label rotamers grafted (*in silico*) on CPR were computed (Figure S8), as well as FMNH^•^-NO^•^ distance distributions (Figure S9), which can then be compared to experimental data. For the position 157, calculated distance distributions were similar in broadening for all 3 labels. For position 668, narrower calculated distance distributions in both states were obtained for the HO-4120 label (cyan line in figure S9). Additionally, no modification in distance distributions was observed between “locked/closed” and “unlocked/open” conformations for Q157^*p*AzPhe*^, as expected in comparison to previously reported results.^[17]^ On the contrary, a dramatic shift in distance was observed for K668^*p*AzPhe*^ between the aforementioned conformations. In both cases, the calculated results were independent of the nature of the label used. Additionally, MMM calculations predicted shorter FMNH^•^-NO^•^ distances (27 Å) for FMNH^•^/K668^*p*AzPhe/14N-HO-5223^ and for FMNH^•^/K668^*p*AzPhe/15N-HO-5223^ than those experimentally measured (31 Å). This difference may arise from a larger separation than predicted between the two domains, or a larger environment occupied by the label at Q157 position, as already hypothesized from the cw EPR data. Despite these variations, the HO-5223 labels were highly effective reporters of domain movements within proteins, as they clearly reflected the impact of ionic strength on the protein’s open conformation. This demonstrated the sensitivity of HO-5223 to conformational changes, making it a valuable tool for studying protein dynamics in solution.

## Conclusion

Conventional SDSL-EPR approaches targeting cysteine residues are limited to proteins where cysteine can be mutated and/or labelled. To address this limitation, we introduced the synthesis and application of novel ncaa/nitroxide pairs designed to extend the capabilities of SDSL-EPR approach for studying protein dynamics. In this work, using acid-catalyzed reaction with the *p*AcPhe/HO-4120 pair and SPAAC reaction with the *p*AzPhe/HO-5223 pair, we successfully labelled the Cytochrome P450 reductase (CPR), a diflavin-containing protein. Expanding the diversity of ncaa/nitroxide pairs is crucial for tailoring chemical approaches to suit specific biological systems and contexts Continuous wave EPR spectroscopy allowed the investigation of the protein dynamics, while DEER measurements confirmed previous findings, notably the presence of two states of CPR in solution in the specific FAD/FMNH^•^ redox state. The introduction of the ^15^N-HO-5223 label proved particularly valuable. Its distinctive two-line EPR spectrum provided same structural insights comparable to those obtained with the ^14^N-isotope, highlighting its utility for studying protein dynamics in solution. The integration of the ^15^N-isotope with ncaa technology presents new perspectives for distances measurements without the need for an additional paramagnetic probe, when using ^14^N- and ^15^N nitroxide simultaneously. Indeed, the use of the ^15^N-label significantly enhances sensitivity, as its signal is distributed over only two lines rather than three, resulting in a more concentrated and detectable response. Further studies could focus on exploring conformational changes of CPR across different redox states (FAD/FMN, FAD/FMNH^•^ and FADH2/FMNH_2_) and under varying ionic conditions, utilizing two incorporated ncAAs instead of relying on the flavin co-factor. Beyond expanding the ncaa/nitroxide pair toolkit for systems where cysteine modification is unfeasible, labels used in this work demonstrated their effectiveness in reporting on protein dynamics and structural changes. Notably, due to the length of their linkers, these labels may also serve as probes for more distant environments. This work marks a significant advancement toward advancing structural studies of flavoproteins, and other proteins. By pushing the boundaries of EPR-based techniques, we pave the way for more versatile and detailed explorations of protein dynamics in complex systems.

## Supporting Information

The authors have cited additional references within the Supporting Information.

## Acknowledgements

This work was supported by the Centre National de la Recherche Scientifique (CNRS) and Aix-Marseille Université (AMU). We are grateful to the EPR-MRS facilities of the Aix-Marseille EPR centre and acknowledge the financial support of the French research infrastructure INFRANALYTICS (FR2054). The authors acknowledge support from MOSBRI which has received funding from the European Union’s Horizon 2020 Research and Innovation Program under Grant Agreement N° 101004806. This project has received funding from National Research, Development, and Innovation Fund of Hungary, NKFI K 137793. We thank Prof. Wayne Hubbell and Evan Brooks, from Jules Stein Eye Institute at UCLA, for helpful discussions on labelling reactions, Prof. Gunnar Jeschke, from ETH Zürich, for implementation of the HO-4451 and HO-5223 labels in MMM software. We thank AMU fellowship for M1 internship for Colette Maignet-Magard. This work was supported by the computing facilities of the CRCMM, ‘Centre Régional de Compétences en Modélisation Moléculaire de Marseille’.

## Notes

### Competing Interest Statement

The authors have declared no competing interest.

