## supplemental information for "Expanding the diversity of nitroxide-based paramagnetic probes conjugated to non-canonical amino acids for SDSL-EPR applications"

[c] Protein Expression Facility

Aix Marseille Univ, CNRS, IMM

13402 Marseille (France)

[d] TBI, Université de Toulouse, CNRS, INRAE, INSA, Toulouse (France)

[e] Szentágothai Research Centre, Ifjúság st. 20, H-7624 Pécs, Hungary

### General information

### Synthesis of $^{14}N/^{15}N$-HO-5223 spin labels

Melting points were determined with a Boetius micro-melting point apparatus and were uncorrected. Elemental analyses (C, H, N, and S) were performed with a Fisons EA 1110 CHNS elemental analyzer. The mass spectra were recorded with a Shimadzu GCMS-2020Q operated in EI mode (70 eV). The $^{1}H$ NMR spectra were recorded with a Bruker Avance 3 Ascend 500 system operated at 500 MHz, and the $^{13}C$ NMR spectra were obtained at 125 MHz in DMSO-$d_6$ at 298 K. The NMR spectra of compound **3** could not be taken by in situ reduction with diphenylhydrazine because of destruction of compound, therefore its diamagnetic O-acetyl derivative was prepared for measurements, as described previously.[1] IR spectra were recorded with a Bruker Alpha FT-IR instrument with ATR support (diamond plate). Flash column chromatography was performed on Merck Kieselgel 60 (0.040–0.063 mm). The TLC was run on plastic 20 × 20 cm 60 $F_{254}$ sheets. Compound **1**[2] and its $^{15}N$ analogue[3] was prepared as described previously; compound **3** was purchased from Merck (Product #: 742678), other reagents were purchased from Merck, Molar, or Novochem.

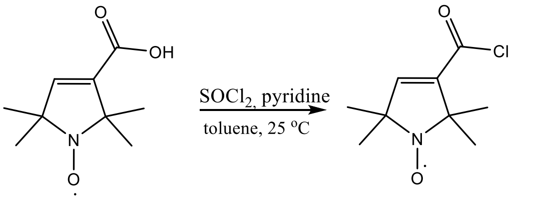

**1** **2**

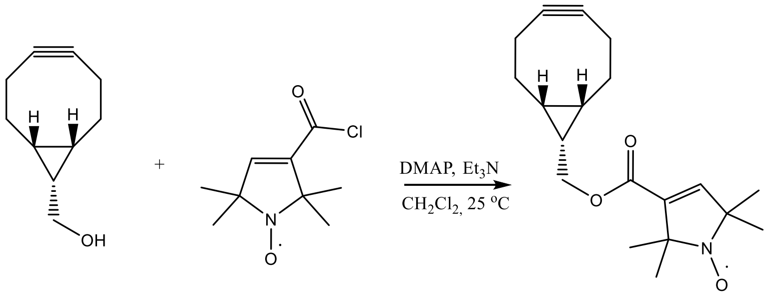

49%

**3** **2** **4**

#### (3-Chloroformyl-2,2,5,5-tetramethyl-2,5-dihydro-1H-pyrrole-1-yl)oxydanyl (**2**)

The paramagnetic acid chloride synthesis was conducted as described earlier with minimal alterations with both $^{14}N$ and 15N pyrroline carboxylic acid as described earlier[1,2]: To the stirred solution of compound **1** (280 mg, 1.5 mmol) and pyridine (0.15 mL, 1.9 mmol) in dry toluene (10 mL), thionyl chloride (0.15 mL, 2.0 mmol) was added dropwise at 0 °C. The mixture was continuously stirred at 25 °C for 1 h. The salts (pyridine/HCl) which precipitated was filtered off, and washed with dry toluene (5 mL), the filtrate was evaporated in a vacuum. Compound **2** was directly used for the synthesis of compound **4**.

##### (((1R,8S,9s)-bicyclo[6.1.0]non-4-yn-9-ylmethyl 2,2,5,5-tetramethyl-2,5-dihydro-1H-pyrrole-3-carboxylate)-1-yl)oxydanyl (**4**)

To a stirred solution of (1R,8S,9s)-bicyclo[6.1.0]non-4-yn-9-ylmethanol (**3**) (80 mg, 0.5 mmol), DMAP (5 mg, 0.04 mmol) and $Et_3N$ (400 mg, 4.0 mmol) in dry $CH_2Cl_2$ (15 mL), paramagnetic acid chloride **2** (202 mg, 1.0 mmol dissolved in 5 mL $CH_2Cl_2$) was added at 0 °C. The reaction mixture was stirred at room temperature overnight. The reaction was quenched with saturated aq. $NH_4Cl$ solution (5 mL), $CH_2Cl_2$ (10 mL) was added and the organic phase was separated, then washed with saturated aq. NaCl solution (10 mL), dried ($MgSO_4$), filtered, evaporated, the crude product was purified by flash chromatography (hexane/$Et_2O$ 2:1); yield: 83 mg (49%); yellow crystal; mp.:110-112 °C; TLC (hexane/EtOAc, 2:1): $R_f$ = 0.47.

IR: 2933, 1714, 1625 $cm^{-1}$.

$^{1}H$ NMR (500 MHz, DMSO-$d_6$ + (N-OAc): δ = 6.70 (s, 1H), 4.24 (d, 2H, J = 8.5Hz), 2.29-2.15 (m, 6H), 2.12 (s, 3H), 1.58 (q, 2H J = 10Hz), 1.36 (d, 1H, J = 8.5 Hz), 1.32 (s, 6H), 1.23 (s, 6H), 0.93 (t, 2H, J = 98Hz).

$^{13}C$ NMR (125 MHz, DMSO-$d_6$ + (N-OAc): δ = 171.1 (1C), 163.3 (1C), 145.9 (1C), 135.9 (1C), 99.4 (2C), 70.4 (1C), 68.7 (1C), 62.6 (1C), 29.1 (1C), 28.3 (1C)., 28.1 (1C), 22.8 (1C), 22.7 (1C), 21.3 (2C), 20.2 (2C), 19.4 (1C), 17.6 (2C).

MS (EI): m / z (%): 316 ([$M^+$], 9), 287 (3), 184 (8), 154 (38), 133 (39), 117 (51), 91 (100).

Anal. Calcd. for $C_{19}H_{26}NO_3$ : C, 72.12; H, 8.28; N, 4.43 Found: C, 72.05 ; H, 8.20; N, 4.39.

##### (((1R,8S,9s)-bicyclo[6.1.0]non-4-yn-9-ylmethyl 2,2,5,5-tetramethyl-2,5-dihydro-1H-pyrrole-3-carboxylate)-1-yl)oxydanyl (**4**) with $^{15}N$ isotope

The same procedure as described above, the resulted compound was a yellow solid with the same physical and chemical properties as for $^{14}N$ analogue.

IR: 2923, 1714, 1629 $cm^{-1}$.

MS (EI): m / z (%): 317 ([$M^+$], 12), 287 (4), 137 (29), 133 (24), 117 (71), 109 (100), 91 (98).

#### CPR sample preparation

##### Expression and purification of soluble forms of Q157$^{pAzPhe/pAcPhe}$ and K668$^{pAzPhe/pAcPhe}$ CPR mutants

All human CPR protein mutants have been truncated on the 66 first amino acids (hereafter named as His-Tag CPR, see below), numbering based on the NCBI Reference Sequence NP_001382342 (UniProt, see below).

```
                   1       10        20        30        40        50        60        70        80        90        100       110       120       130
                   +--------+---------+---------+---------+---------+---------+---------+---------+---------+---------+---------+---------+---------+
UniProt          : MGDSHVDTSSTVSEAVAEEVSLFSMTDMILFSLIVGLLTYWFLFRKKKEEVPEFTKIQTLTSSVRESSFVEKMKKTGRNIIVFYGSQTGTAEEFANRLSKDAHRYGMRGMSADPEEYDLADLSSLPEIDN
His-Tag CPR      : --------------------------------------MGSSHHHHHHSSGLVPRGSHMLESSVRESSFVEKMKKTGRNIIVFYGSQTGTAEEFANRLSKDAHRYGMRGMSADPEEYDLADLSSLPEIDN
CPR (PDB 3QE2)   : -----------------------------------------------------------------ESSFVEKMKKTGRNIIVFYGSQTGTAEEFANRLSKDAHRYGMRGMSADPEEYDLADLSSLPEIDN

                            140       150       160       170       180       190       200       210       220       230       240       250       260
                   ---------+---------+---------+---------+---------+---------+---------+---------+---------+---------+---------+---------+---------+
UniProt          : ALVVFCMATYGEGDPTDNAQDFYDWLQETDVDLSGVKFAVFGLGNKTYEHFNAMGKYVDKRLEQLGAQRIFELGLGDDDGNLEEDFITWREQFWPAVCEHFGVEATGEESSIRQYELVVHTDIDAAKVYM
His-Tag CPR      : ALVVFCMATYGEGDPTDNAQDFYDWLQETDVDLSGVKFAVFGLGNKTYEHFNAMGKYVDKRLEQLGAQRIFELGLGDDDGNLEEDFITWREQFWPAVCEHFGVEATGEESSIRQYELVVHTDIDAAKVYM
CPR (PDB 3QE2)   : ALVVFCMATYGEGDPTDNAQDFYDWLQETDVDLSGVKFAVFGLGNKTYEHFNAMGKYVDKRLEQLGAQRIFELGLGDDDGNLEEDFITWREQFWLAVCEHFGVEAT---SSIRQYELVVHTDIDAAKVYM

                            270       280       290       300       310       320       330       340       350       360       370       380       390
                   ---------+---------+---------+---------+---------+---------+---------+---------+---------+---------+---------+---------+---------+
UniProt          : GEMGRLKSYENQKPPFDAKNPFLAAVTTNRKLNQGTERHLMHLELDISDSKIRYESGDHVAVYPANDSALVNQLGKILGADLDVVMSLNNLDEESNKKHPFPCPTSYRTALTYYLDITNPPRTNVLYELA
His-Tag CPR      : GEMGRLKSYENQKPPFDAKNPFLAAVTTNRKLNQGTERHLMHLELDISDSKIRYESGDHVAVYPANDSALVNQLGKILGADLDVVMSLNNLDEESNKKHPFPCPTSYRTALTYYLDITNPPRTNVLYELA
CPR (PDB 3QE2)   : GEMGRLKSYENQKPPFDAKNPFLAAVTTNRKLNQGTERHLMHLELDISDSKIRYESGDHVAVYPANDSALVNQLGKILGADLDVVMSLNNLDEESNKKHPFPCPTSYRTALTYYLDITNPPRTNVLYELA

                            400       410       420       430       440       450       460       470       480       490       500       510       520
                   ---------+---------+---------+---------+---------+---------+---------+---------+---------+---------+---------+---------+---------+
UniProt          : QYASEPSEQELLRKMASSSGEGKELYLSWVVEARRHILAILQDCPSLRPPIDHLCELLPRLQARYYSIASSSKVHPNSVHICAVVVEYETKAGRINKGVATNWLRAKEPAGENGGRALVPMFVRKSQFRL
His-Tag CPR      : QYASEPSEQELLRKMASSSGEGKELYLSWVVEARRHILAILQDCPSLRPPIDHLCELLPRLQARYYSIASSSKVHPNSVHICAVVVEYETKAGRINKGVATNWLRAKEPAGENGGRALVPMFVRKSQFRL
CPR (PDB 3QE2)   : QYASEPSEQELLRKMASSSGEGKELYLSWVVEARRHILAILQDCPSLRPPIDHLCELLPRLQARYYSIASSSKVHPNSVHICAVVVEYETKAGRINKGVATNWLRAKEPV-----RALVPMFVRKSQFRL

                            530       540       550       560       570       580       590       600       610       620       630       640       650
                   ---------+---------+---------+---------+---------+---------+---------+---------+---------+---------+---------+---------+---------+
UniProt          : PFKATTPVIMVGPGTGVAPFIGFIQERAWLRQQGKEVGETLLYYGCRRSDEDYLYREELAQFHRDGALTQLNVAFSREQSHKVYVQHLLKQDREHLWKLIEGGAHIYVCGDARNMARDVQNTFYDIVAEL
His-Tag CPR      : PFKATTPVIMVGPGTGVAPFIGFIQERAWLRQQGKEVGETLLYYGCRRSDEDYLYREELAQFHRDGALTQLNVAFSREQSHKVYVQHLLKQDREHLWKLIEGGAHIYVCGDARNMARDVQNTFYDIVAEL
CPR (PDB 3QE2)   : PFKATTPVIMVGPGTGVAPFIGFIQERAWLRQQGKEVGETLLYYGCRRSDEDYLYREELAQFHRDGALTQLNVAFSREQSHKVYVQHLLKQDREHLWKLIEGGAHIYVCGDARNMARDVQNTFYDIVAEL

                            660       670
                   ---------+---------+-------
UniProt          : GAMEHAQAVDYIKKLMTKGRYSLDVWS
His-Tag CPR      : GAMEHAQAVDYIKKLMTKGRYSLDVWS
CPR (PDB 3QE2)   : GAMEHAQAVDYIKKLMTKGRYSLDVWS
```

Production of Q157$^{p\text{AzPhe}}$, K668$^{p\text{AzPhe}}$, Q157$^{p\text{AcPhe}}$ and K668$^{p\text{AcPhe}}$ CPR mutants was performed as previously described.[4] BL21 DE3 strains were provided from New England BioLabs (NEB, catalogue number C2527I). A 10 mL pre-culture was inoculated from a single colony glycerol stock, and grown overnight at 37°C in LB medium supplemented with 100 µg/mL Ampicilline and 34 µg/mL Cloramphenicol. In an Erlenmeyer of 1L, 2 mL of the grown pre-culture was used to inoculate, at 28°C for 5h, 200 mL TB medium supplemented with 100 µg/mL Ampicilline, 34 µg/mL Cloramphenicol, 1 mM $MgSO_4$, 100 µM Riboflavin, 50 mg *p*AzPhe or 50 mg *p*AcPhe. The production of Q157$^{p\text{AzPhe}}$ and K668$^{p\text{AzPhe}}$ is carried out into darkness to avoid the degradation of *p*AzPhe ncaa. In order to obtain a soluble, well folded and functional CPR, the culture was grown without addition of IPTG to took advantage of the leakage of the *lac* operon and. 400 µL of 10 % L-Arabinose (0.02 % final concentration) was added and the culture was grown for additional 43h. Cells were harvested by centrifugation (7000 x g, 15min, 4°C). Cell pellets were resuspended and homogenized using a Potter in lysis buffer (20 mM Tris HCl pH 7.4, 250 mM NaCl, 250 mM KCl, 1 x protease inhibitor cocktail [aprotinin (0.3 µM), leupeptin (1 µM), pepstatin A (1.5 µM), benzamidine (100 µM), sodium metabisulfite (100 µM)] Lysozyme and DNase I). The cells were disrupted 2 times through the cell disruptor (CellD, 1.8kBar), then centrifuged (1h, 12 000 x g). The kept supernatant was loaded on a Hitrap TALON crude 5mL column (GE Healthcare), washed (20 mM Tris HCl buffer pH 7.4, 250 mM NaCl, 250 mM KCl, 10 mM imidazole) and eluted with a containing imidazole buffer (20 mM Tris HCl buffer pH 7.4, 250 mM NaCl, 250 mM KCl, 300 mM imidazole). The protein-containing fractions were recovered and loaded on a 2'5'-ADP- Sepharose column (GE Healthcare), washed (20 mM Tris HCl buffer pH 7.4, 150 mM NaCl) and eluted with a $NADP^+$ buffer containing (20 mM Tris HCl buffer pH 7.4, 150 mM NaCl, 1 mM $NADP^+$). To better quantify the CPR amount, the purified protein was fully oxidized with a high excess amount of potassium ferricyanide (100 mM stock solution) supplemented with an excess of FAD and FMD. Excess of oxidant and of flavines was removed by ultrafiltration using a 30 kDa cut-off polyethersulfonemembrane (PES) (Vivaspin, Sartorius). Amount of soluble CPR was quantified by absorbance at 455 nm (Epsilon = 21 500 $cm^{-1}.M^{-1}$).

#### SDS-PAGE gels

Protein integrity was verified using a 12% SDS-PAGE: placing samples (mix with loading buffer containing SDS, DTT, 2-betamercaptoethanol and heat at 96°C for 5 min) in gel wells, running for 15 min at 90 V then 1 hour at 120 V in running buffer 1X (stock 10X: Tris 30g.$L^{-1}$ glycine 80 g.$L^{-1}$ and SDS 5 g.$L^{-1}$). For the sake of clarity, we only kept gel lines (of fractions containing the protein after purification on TALON and 2’5’-ADP-Sepharose columns: total fraction, soluble part, flow-through, elution. Lanes from the same gel have been spliced together (indicated by vertical lines in figure S3 and S4).

#### Activity assays

Activity assays were based on the initial velocity ($k_{obs}$) of reduction of cytochrome *c* by CPR in presence of NADPH, by monitoring the increase of the absorption band at 550nm using a Cary 60 Scan UV-Vis spectrophotometer (Agilent, France). The mixture was prepared in Tris 20 mM buffer pH 7.4, in a plastic cuvette (1 cm light path, 1 mL total volume): put 943 µL of buffer (record the blank at 550 nm), start the kinetics measurement, add quickly 50 µL cytochrome *c* from bovine heart (2 mM stock solution, Sigma-Aldrich) and mix by inverting the cuvette. Record for few seconds the signal of the cytochrome *c* alone. Add quickly 2 µL of NADPH (100 mM frozen stock solution, Carbosynth), mix again by inverting the cuvette and verify that NADPH does not reduce the cytochrome *c*. Finally, add 5 µL of CPR (0.5 µM stock concentration, supplemented with an excess of 0.2 µM FAD and 2 µM FMN to avoid the loss of the cofactors). Then record the kinetics for 10 minutes. The initial velocity $k_{obs}$ was calculated using the slope of the linear part of the curve, using the molar extinction coefficient of Cytochrome *c* (21 000 $cm^{-1}.M^{-1}$) and the concentration of CPR in the cuvette (2.5 nM).[4] For each activity assay, an experimental triplicate were done. Error bars were calculated using Origin software.

#### Labelling reaction

##### *Labelling protocol using the pAzPhe ncaa*

Labelling reaction of $Q157^{pAzPhe}$ or $K668^{pAzPhe}$ was done thanks to a SPAAC click chemistry reaction[5] using the HO-4451, HO-5223 labels.[6] 10-molar fold excess of HO-4451 or HO-5223 (10 mM stock solution in DMSO) was added to a concentrated solution of mutant CPR (200-400 µM final concentration) in Tris 20 mM buffer pH 7.4. The mixture was let to incubate overnight at room temperature, into darkness. Excess of label was removed by using a Centripure P2 Zetadex Gel filtration column (Clinisciences - NeoBiotech) in Tris 20 mM buffer pH 7.4. Fractions containing spin labelled protein (see below) were pooled and concentrated. Labelling yields were calculated by dividing the spin concentration (evaluated by double integration of the EPR signal recorded under non-saturating conditions, see below) by the protein concentration of the pool: in the range of 44 to 53% for $Q157^{pAzPhe}$ and $K668^{pAzPhe}$ respectively, although the yield gets higher (63%) in presence of 500 mM of NaCl for $K668^{pAzPhe}$. Labelled proteins were named hereafter as $Q157^{pAzPhe*}$ or $K668^{pAzPhe*}$ (star indicating a labelled protein). The spin-labelled enzymes were stored at -80°C.

***Labelling protocol using the pAcPhe ncaa***

Labelling reaction of Q157$^{pAcPhe}$ or K668$^{pAcPhe}$ was performed thanks to an oxime reaction using the HO-4120 label. After purification, the protein buffer was exchanged to a sodium phosphate buffer (20 mM), pH 6, NaCl 500 mM. Under argon flux, 0.5 molar fold of catalyst (glycine or *para*-methoxyalanine, 10 mM stock) was added to a concentrated solution of mutant CPR (200-400 µM final concentration). After five minutes, 10-molar fold excess of HO-4120 (10 mM stock solution in DMSO) was added. The mixture was let to incubate 48 to 72h at room temperature, into darkness. Excess of label was removed using a Centripure P2 Zetadex Gel filtration column (Clinisciences - NeoBiotech) in Tris 20 mM buffer pH 7.4. Fractions containing spin labelled protein were pooled and concentrated. Labelling yields were calculated by dividing the spin concentration (evaluated by double integration of the EPR signal recorded under non-saturating conditions, see below) by the protein concentration of the pool : in the range of 58 to 81% for Q157$^{pAcPhe}$ and K668$^{pAcPhe}$ respectively. Labelled proteins were named hereafter as Q157$^{pAcPhe*}$ or K668$^{pAccPhe*}$ (star indicating a labelled protein).

##### Preparation of FMNH$^{\bullet}$ species

1 eq of NADPH (100 mM frozen stock solution, Carbosynth) was added to a concentrated labelled protein Q157$^{pAzPhe*/pAcPhe*}$ or K668$^{pAzPhe*/pAcPhe*}$ in Tris 20 mM buffer pH 7.4. The sample was let to evolve to reach a maximum of FMNH$^{\bullet}$ species as monitored by the 620 nm absorption band increase. The dark green colour of the solution was an indication of the semiquinone species. The FMNH$^{\bullet}$ state is the air stable species obtained after purification of soluble CPR as well.

##### Preparation of DEER samples

To achieve better sensitivity and higher-quality distance distribution measurements, the proteins were placed in Tris 20 mM buffer pH 7.4 prepared in $D_2O$.[7] The final protein concentrations ranged from 80 to 100 µm. $D_8$-glycerol (30% v/v) was added to the samples before rapid freezing to avoid heterogeneous protein concentration.

#### Spectroscopy

##### Continuous wave (CW) EPR spectroscopy

EPR spectra were recorded at room temperature on an ELEXSYS E500II Bruker spectrometer equipped with an ELEXSYS Super High Q sensitivity resonator operating at 9.9 GHz. The magnetic field modulation frequency was 100 kHz. Relaxation parameters of FMNH$^{\bullet}$ and nitroxide labels are quite different at room temperature. Depending on the paramagnetic species (FMNH$^{\bullet}$, labelled protein -CPR*-, or both), the optimised non saturating parameters are given in the table below. The concentration of labelled proteins was evaluated by double integration of the EPR signal recorded under non-saturating conditions and compared with a MTSL standard sample of known concentration.

| | Power (mW) | Modulation Amplitude (mT) |
|---|---|---|
| **FMNH$^{\bullet}$** | 4 | 0.8 |
| **CPR*** | 10 | 0.1 |
| **FMNH$^{\bullet}$/CPR*** | 4 | 0.1 |

##### Pulsed (DEER) spectroscopy

Q-band pulsed EPR experiments were performed on an Elexsys E580 spectrometer (Bruker) using an EN 5107D2 resonator. Temperature was reached with a closed cycle He Stinger system from Bruker and controlled by a Mercury controller from Oxford. The EPR spectrometer was used exploiting an Arbitrary Waveform Generator (AWG) and 10W microwaves solid-state amplifier. The data were recorded at a temperature of 60 K for the DEER experiments. For nitroxide–FMNH$^{\bullet}$ distance measurements, a four-pulse DEER sequence[8] was used with pulse durations of 20 ns (π/2 pulse) and 40 ns (π pulse) with delays $t_1$ of 200 ns and $t_2$ adjusted according to Tm (phase memory time). The pump frequency was set to the maximum of the resonance for the FMNH$^{\bullet}$ radical, and the observed frequency was 70 MHz lower, corresponding to the maximum of the nitroxide spectrum. To suppress unwanted echoes, an eight-step phase cycle was applied. The EFS spectral shape obtained using the $^{15}$N isotope in $^{15}$N-HO-5223 was modified due to the $^{15}$N-hyperfine coupling. Data were collected for about 3-4 h for each sample. Distance distributions obtained using the auto-computed mode in DeerAnalysis 2022.[9,10] Color coding for reliability ranges are : green: Shape of distance distribution is reliable ; yellow: Mean distance and width are reliable ; orange: Mean distance is reliable ; red: Long-range distance contributions may be detectable, but cannot be quantified.

#### Computational Modelling

Each structural model of a marker grafted onto an amino acid was constructed using the AGUI software. An initial geometry optimization step was performed with the Gaussian16 software, employing density functional theory (DFT) with the hybrid functional B3LYP and the 6-31g(d,p) basis set. In a second step, a simulated annealing protocol was applied to the optimized structure to explore the conformational space of the molecule and obtain the various possible conformers. This step was carried out using the AMPAC software with the semi-empirical quantum chemistry method AM1.

#### Supplementary figures and tables

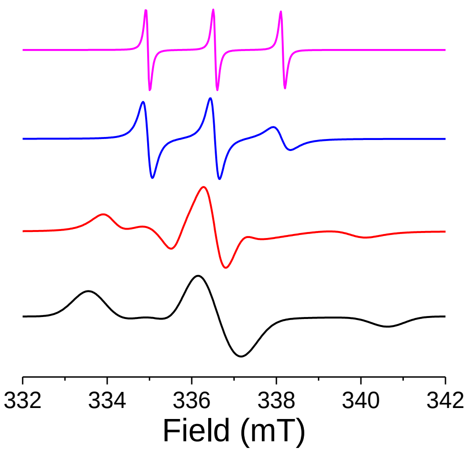

**Figure S1.** Cw EPR spectral shape modifications as a function of the mobility of the spin label $^{14}$N-HO-5223 described by its rotational correlation time $\tau_c$. The spectra have been simulated using EasySpin[11] for different values of $\tau_c$: 0.01 ns (pink spectrum), 1 ns (blue spectrum), 10 ns (red spectrum) and 100 ns (black spectrum).

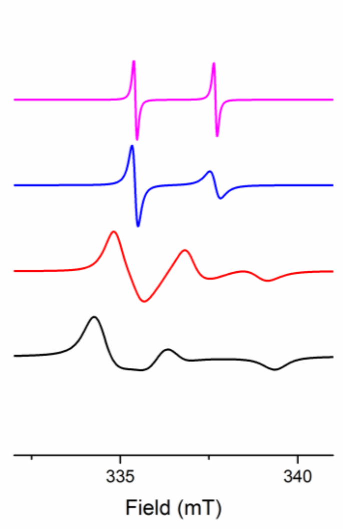

**Figure S2.** Cw EPR spectral shape modifications as a function of the mobility of the spin label $^{15}$N-HO-5223 described by its rotational correlation time $\tau_c$. The spectra have been simulated using EasySpin for different values of $\tau_c$: 0.01 ns (pink spectrum), 1 ns (blue spectrum), 10 ns (red spectrum) and 100 ns (black spectrum).

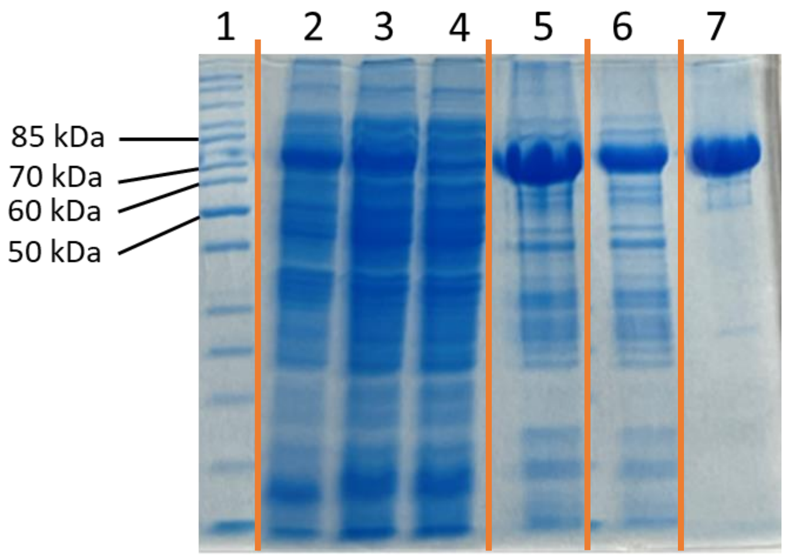

**Figure S3.** 12 % SDS-PAGE of expression Q157$^{pAzPhe}$ CPR mutant. 1: Protein ladder (EZ-Run Rec Protein Ladder, Fisher BioReagents), 2: total fraction, 3: Soluble part, 4: Flow-through (TALON), 5: Elution (TALON), 6: Flow-through(2'5'-ADP-Sepharose), 7: Elution (2'5'-ADP-Sepharose). Running buffer 1X (stock 10X: Tris 30 g.L$^{-1}$ glycine 80 g.L$^{-1}$ and SDS 5 g.L$^{-1}$). Orange vertical lines indicated gel splicing.

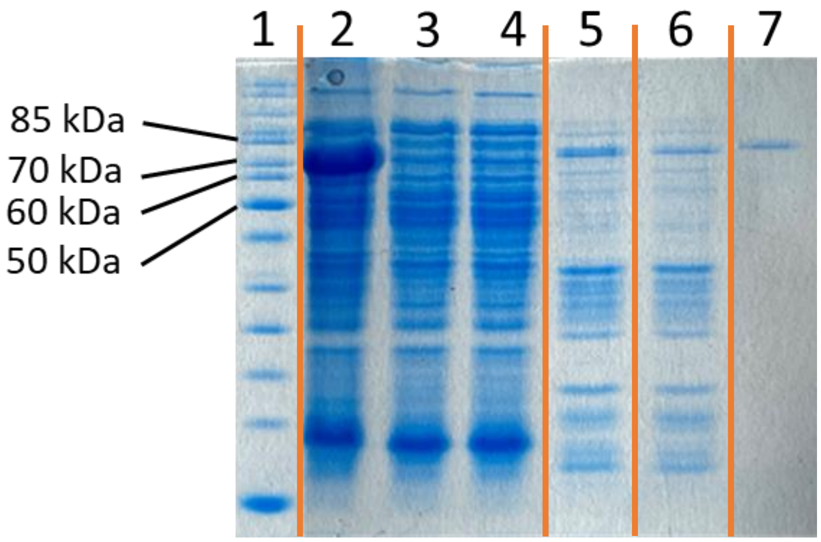

**Figure S4.** 12 % SDS-PAGE gel of expression K668$^{pAzPhe}$ CPR mutant. 1: Protein ladder (EZ-Run Rec Protein Ladder, Fisher BioReagents), 2: total fraction, 3: Soluble part, 4: Flow-through (TALON), 5: Elution (TALON), 6: Flow-through (2'5'-ADP-Sepharose), 7: Elution (2'5'-ADP-Sepharose). Running buffer 1X (stock 10X: Tris 30 g.L$^{-1}$ glycine 80 g.L$^{-1}$ and SDS 5 g.L$^{-1}$). Orange vertical lines indicated gel splicing.

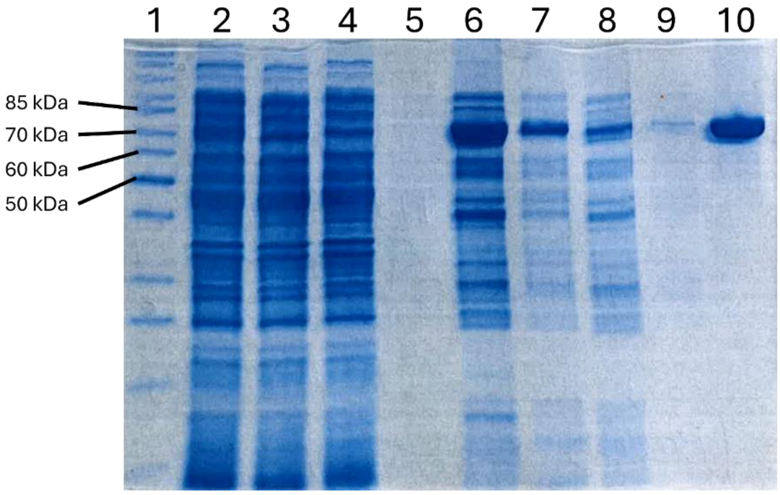

**Figure S5.** 12 % SDS-PAGE gel of expression Q157$^{pAcPhe}$ CPR mutant. 1: Protein ladder (EZ-Run Rec Protein Ladder, Fisher BioReagents), 2: total fraction, 3: Soluble part, 4: Flow-through (TALON), 5: Wash (TALON), 6: Elution fraction 1 (TALON), 7: Elution fraction 2 (TALON), 8: Flow-through (2’5’-ADP-Sepharose), 9: Wash (2’5’-ADP-Sepharose), 10: Elution (2’5’-ADP-Sepharose). Running buffer 1X (stock 10X: Tris 30 g.L$^{-1}$ glycine 80 g.L$^{-1}$ and SDS 5 g.L$^{-1}$).

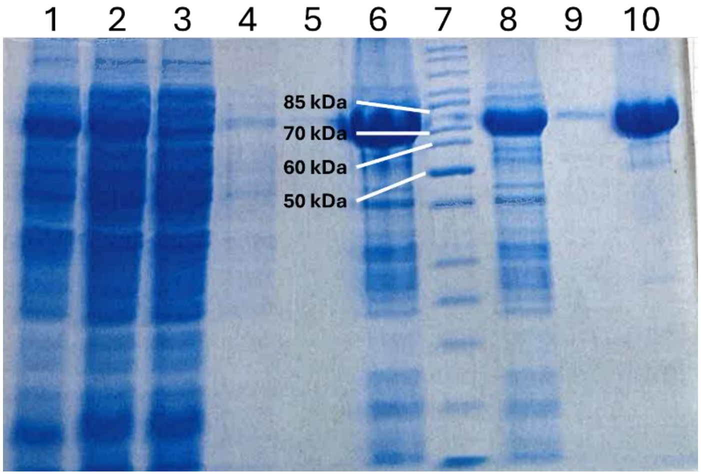

**Figure S6.** 12 % SDS-PAGE gel of expression K668$^{pAcPhe}$ CPR mutant. 1: total fraction, 2: Soluble part, 3: Flow-through (TALON), 4: Wash fraction 1 (TALON), 5: Wash fraction 2 (TALON), 6: Elution (TALON), 7: Protein ladder (EZ-Run Rec Protein Ladder, Fisher BioReagents), 8: Flow-through (2’5’-ADP-Sepharose), 9: Wash (2’5’-ADP-Sepharose), 10: Elution (2’5’-ADP-Sepharose). Running buffer 1X (stock 10X: Tris 30 g.L$^{-1}$ glycine 80 g.L$^{-1}$ and SDS 5 g.L$^{-1}$).

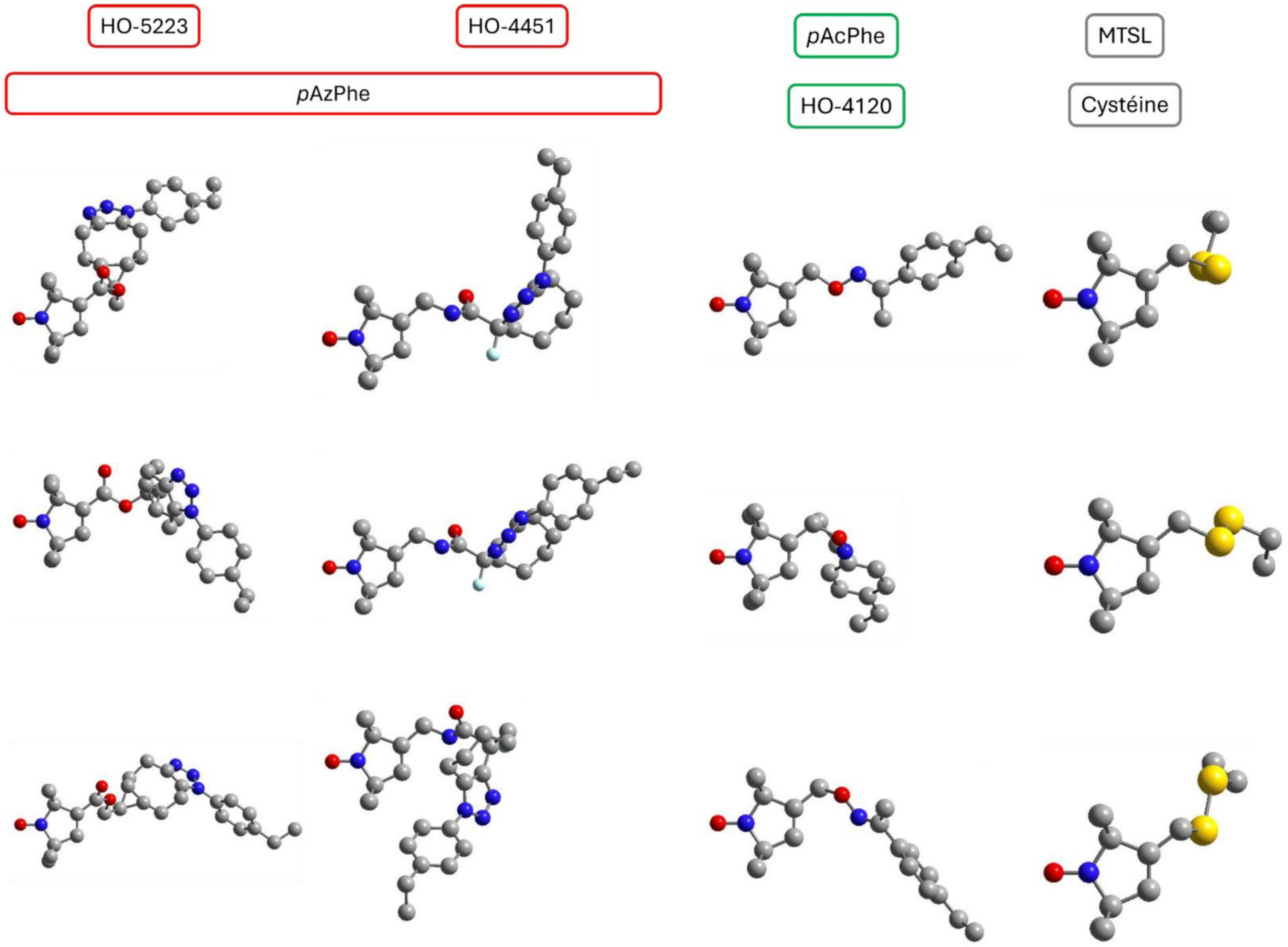
HO-5223
HO-4451
pAzPhe
pAcPhe
HO-4120
MTSL
Cystéine

**Figure S7.** Example of three conformers picked from the set of labels calculated conformers, with the associated ncaa indicated on the left.

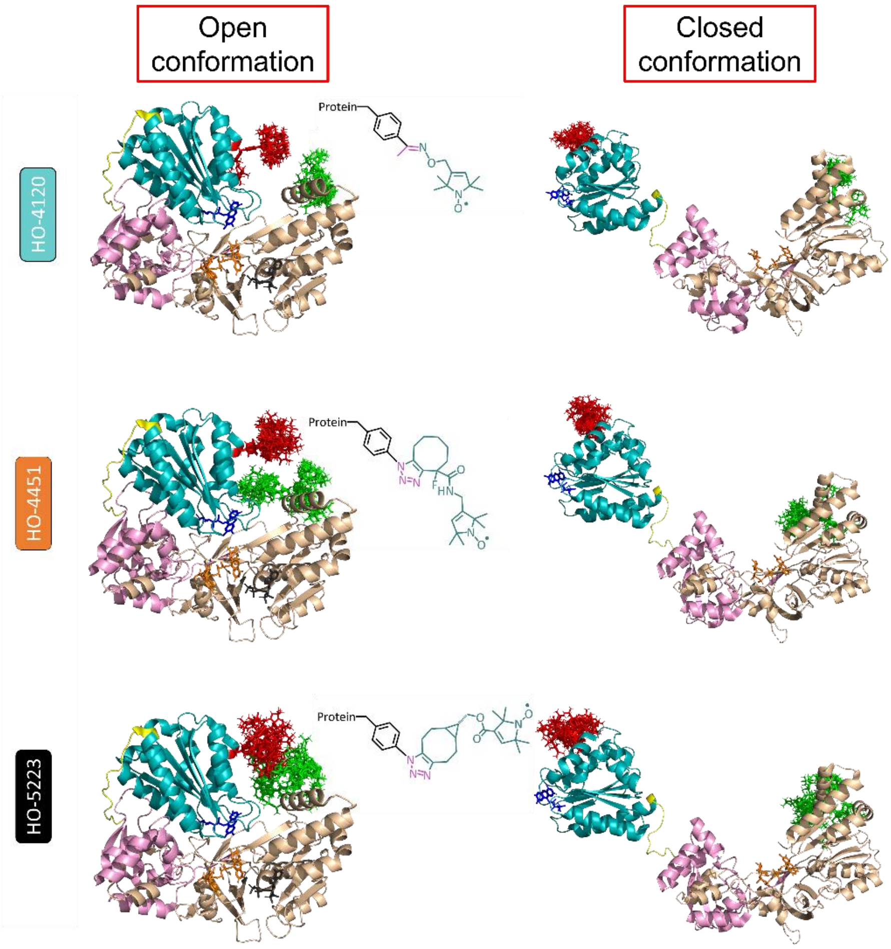

**Figure S8.** MMM[12] computed spin-labelled rotamers with *p*AzPhe attached to positions Q157 (red) and K668 (green) based on the crystal structure of soluble *Homo sapiens* CPR for the “locked/compact” (pdb. 5FA6) conformation and on the *Saccharomyces cerevisiae/Homo sapiens* CPR for the “unlocked/open” (pdb. 3FJO) chimeric conformation. The FAD and FMN domains are represented in clear brown and cyan ribbons respectively, FAD and FMN co-factors are shown as ticks in orange and dark blue, linker is shown as ribbon in light pink, the flexible loop is shown in yellow.

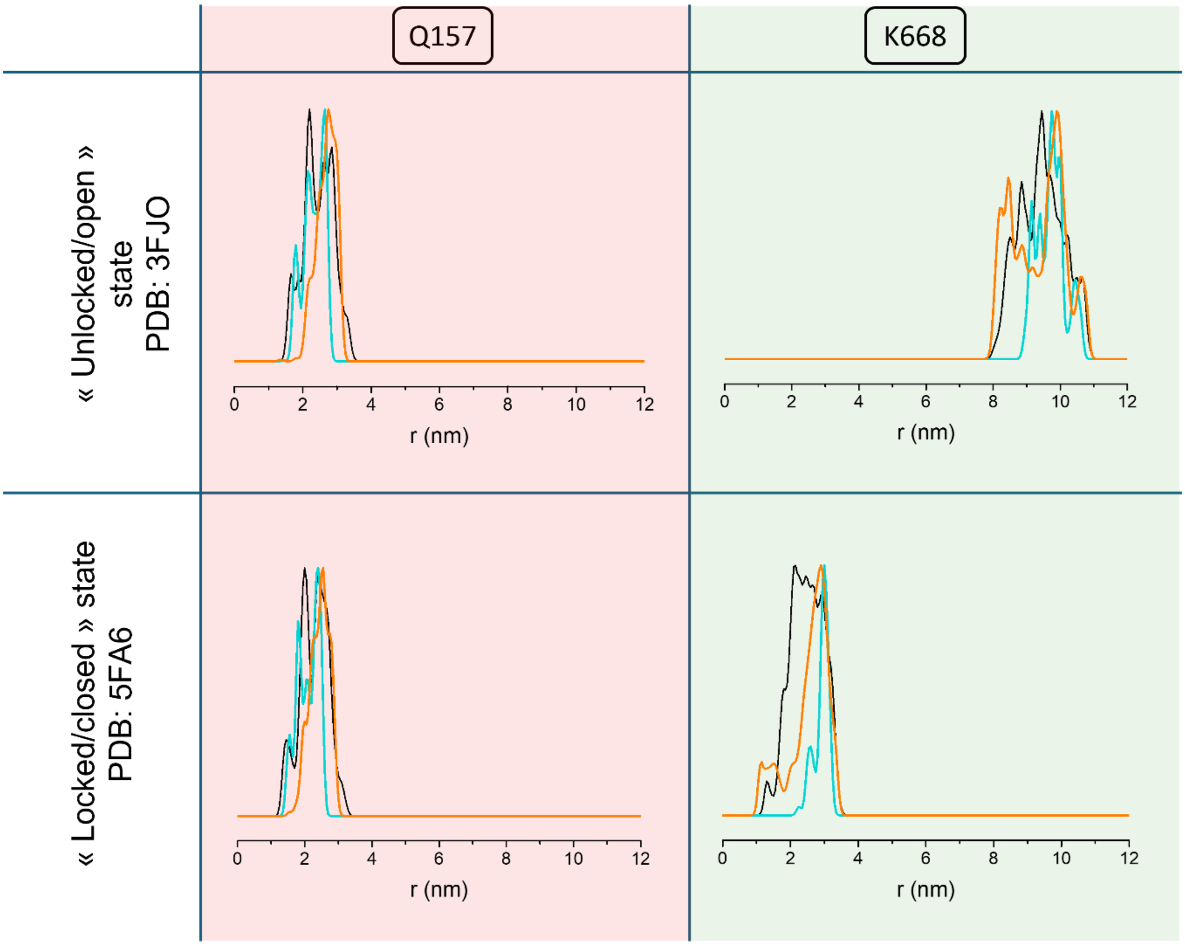

**Figure S9.** MMM[12] computed distance distributions between FMN N5 and nitroxide labels HO-4451 (orange) , HO-5223 (black), HO-4120 (cyan), for positions Q157 (red rectangle), K668 (green rectangle), based on the crystal structure of soluble *Homo sapiens* CPR for the "locked/closed" (pdb. 5FA6) conformation and on the *Saccharomyces cerevisiae/Homo sapiens* CPR for the "unlocked/open" (pdb. 3FJO) chimeric conformation.

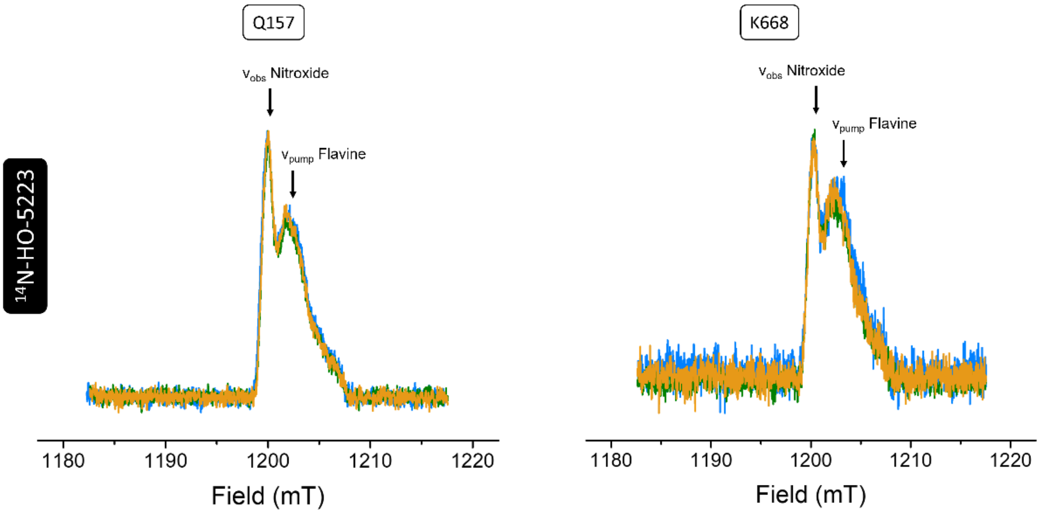

**Figure S10.** Superposition of echo field sweeps (EFS) for Q-band DEER experiments recorded at 60 K on FMNH$^{\bullet}$/Q157$^{pAzPhe/14N\text{-}HO\text{-}5223}$ (left) and FMNH$^{\bullet}$/K668$^{pAzPhe/14N\text{-}HO\text{-}5223}$ (right) in absence (blue line) or in presence of 50 mM (green line), 250 mM (orange line) of NaCl. [K668$^{pAzPhe*}$] = 90 µM, [Q157$^{pAzPhe*}$] = 90 µM. Buffer: Tris HCl 20 mM pH 7.4, $D_2O$, 10% v/v $d_8$-glycerol. Shot repetition time (srt) = 2000 ns.

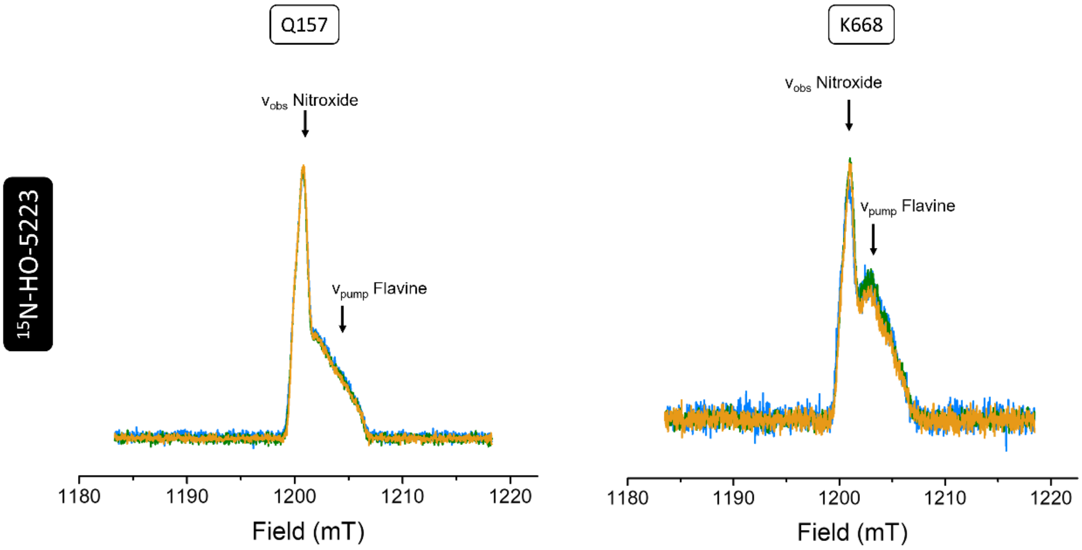

**Figure S11.** Superposition of echo field sweeps (EFS) for Q-band DEER experiments recorded at 60 K on FMNH$^{\bullet}$/Q157$^{pAzPhe/15N\text{-}HO\text{-}5223}$ (left) and FMNH$^{\bullet}$/K668$^{pAzPhe/15N\text{-}HO\text{-}5223}$ (right) in absence (blue line) or in presence of 50 mM (green line), 250 mM (orange line) of NaCl. [K668$^{pAzPhe*}$] = 90 µM, [Q157$^{pAzPhe*}$] = 90 µM. Buffer: Tris HCl 20 mM pH 7.4, $D_2O$, 10% v/v $d_8$-glycerol. Shot repetition time (srt) = 2000 ns. The EFS spectral shape obtained using the $^{15}N$ isotope in $^{15}N$-HO-5223 was modified due to the $^{15}N$-hyperfine coupling.

**Table S1.** Results of CPR production under different conditions.

| Ncaa | Sample | Medium (Culture time) | T (°C) | Volume of culture (mL) | Volume of pure protein (µL) | Concentration (µM) | Quantity (mg) |
|---|---|---|---|---|---|---|---|
| *p*AcPhe | CPR Q157$^{pAcPhe}$ | TB (48h) | 28 | 400 | 600 | 120 | 10 |
| | CPR K668$^{pAcPhe}$ | TB (48h) | 28 | 400 | 1200 | 121 | 10,5 |
| *p*AzPhe | CPR Q157$^{pAzPhe}$ (Darkness) | TB (48h) | 28 | 400 | 580 | 42 | 3,6 |
| | CPR K668$^{pAzPhe}$ (Darkness) | TB (48h) | 28 | 400 | 400 | 448 | 14 |

**Table S2.** Summary of CPR Q157$^{pAcPhe}$, Q157$^{pAcPhe/HO\text{-}4120}$ and WT activities.

| Sample | CPR WT | CPR Q157$^{pAcPhe}$ | CPR Q157$^{pAcPhe/4120}$ | CPR Q157$^{pAcPhe}$ without nitroxide |
|---|---|---|---|---|
| **Activity** | 2760 min$^{-1}$ | 2569 min$^{-1}$ | 1120 min$^{-1}$ | 1325 min$^{-1}$ |
